# A global LC-MS^2^-based methodology to identify and quantify anionic phospholipids in plant samples

**DOI:** 10.1101/2023.05.10.539967

**Authors:** Manon Genva, Louise Fougère, Delphine Bahammou, Sébastien Mongrand, Yohann Boutté, Laetitia Fouillen

## Abstract

Anionic phospholipids (PS, PA, PI, PIPs) are low abundant phospholipids with impactful functions in cell signaling, membrane trafficking and cell differentiation processes. They can be quickly metabolized and can transiently accumulate at define spots within the cell or an organ to respond to physiological or environmental stimuli. As even a small change in their composition profile will produce a significant effect on biological processes, it is crucial to develop a sensitive and optimized analytical method to accurately detect and quantify them. While thin layer chromatography (TLC) separation coupled with gas chromatography (GC) detection methods already exist, they do not allow for precise, sensitive and accurate quantification of all anionic phospholipid species. Here we developed a method based on high performance liquid chromatography (HPLC) combined with two-dimensional mass spectrometry (MS^2^) by MRM mode to detect and quantify all molecular species and classes of anionic phospholipids in one-shot. This method is based on a derivatization step by methylation that greatly enhances the ionization, the separation of each peaks, the peak resolution as well as the limit of detection and quantification for each individual molecular species, and more particularly for PA and PS. Our method universally works in various plant samples. Remarkably, we identified that PS is enriched with very long chain fatty acids in the roots but not in aerial organs of *Arabidopsis thaliana*. Our work thus paves the way to new studies on how the composition of anionic lipids is finely tuned during plant development and environmental responses.

**Significance Statement:** While anionic phospholipids have key functions in plant cellular processes, their low concentration in biological samples and their low stability during the analysis complicate their quantification. Here, we present the first one-shot analytical method for the profiling and quantification of all anionic phospholipid classes and species from plant tissues with unprecedented sensitivity. This method open the way to future studies requiring a fine quantification of anionic phospholipids to understand their role in plant cell processes.

## 1. Introduction

Lipids are not only essential building blocks of cellular membranes but they also play key regulatory functions in a high variety of biological processes (Cai *et al*., 2016; Noack and Jaillais, 2020; Dubois and Jaillais, 2021). Among these, the role of lipids in cellular processes is receiving more and more attention, especially in organisms like plants that should quickly adapt to environmental changes (Wang *et al*., 2016; Noack and Jaillais, 2020; Boutté and Jaillais, 2020). Plant lipids display a high structural diversity of molecules amongst which anionic phospholipids, despite being low abundant, are widely recognized for their role in most membrane-related processes including cell growth, cell division, signaling, membrane trafficking, cell polarity or stress response (Noack and Jaillais, 2020; Dubois and Jaillais, 2021; Noack and Jaillais, 2017; Platre *et al*., 2019; Pokotylo *et al*., 2018). Lipids modulate the physicochemical properties of plant membranes, notably their charge, curvature and the clustering of specific lipids and proteins into membrane domains (Noack and Jaillais, 2020). In addition, anionic phospholipids establish a membrane lipid identity to precisely target membrane processes by allowing the recruitment of specific proteins or effectors such as small GTPases of the ADP ribosylation factor (ARF), the Rab or the Rho families or their activators (the guanine nucleotide exchange factors GEF) or inactivators (the GTPase activating protein GAP) (Noack and Jaillais, 2020; Dubois and Jaillais, 2021; Noack and Jaillais, 2017; Platre *et al*., 2019; Naramoto *et al*., 2009). Additionally, anionic phospholipids are key molecules in cell signaling, which is enabled by their fast biosynthesis, degradation or consumption to produce secondary messengers (Noack and Jaillais, 2020). Accumulation of anionic phospholipid species at specific sub-domains of membrane compartments is tightly regulated by enzymes that either produce or degrade them, including kinases, phosphatases and phospholipases. Thus, the localization or the accumulation of a particular anionic phospholipid specie can be only transient before it gets metabolized into another specie. Anionic phospholipids (**Figure 1**) contain a negatively charged polar head and include the phosphatidic acid (PA), which plays a central role in the biosynthesis of all phospholipid species in plants (Dubois and Jaillais, 2021; Dubots *et al*., 2012), as well as phosphatidylserine (PS), phosphatidylinositol (PI) and their phosphorylated forms called the phosphoinositides. The latter comprises in plants mono- and bis-phosphorylated forms and are respectively named as monophosphate phosphoinositides (PIPs) and bisphosphate phosphoinositides (PIP_2_) (Dubois and Jaillais, 2021). Of note, while phosphatidylinositol-3,4,5-triphosphate (PtdIns(3,4,5)P_3_) is described in animal cells, it is apparently absent in plants (Heilmann and Heilmann, 2015; Molnár *et al*., 2016).

**Figure 1.**
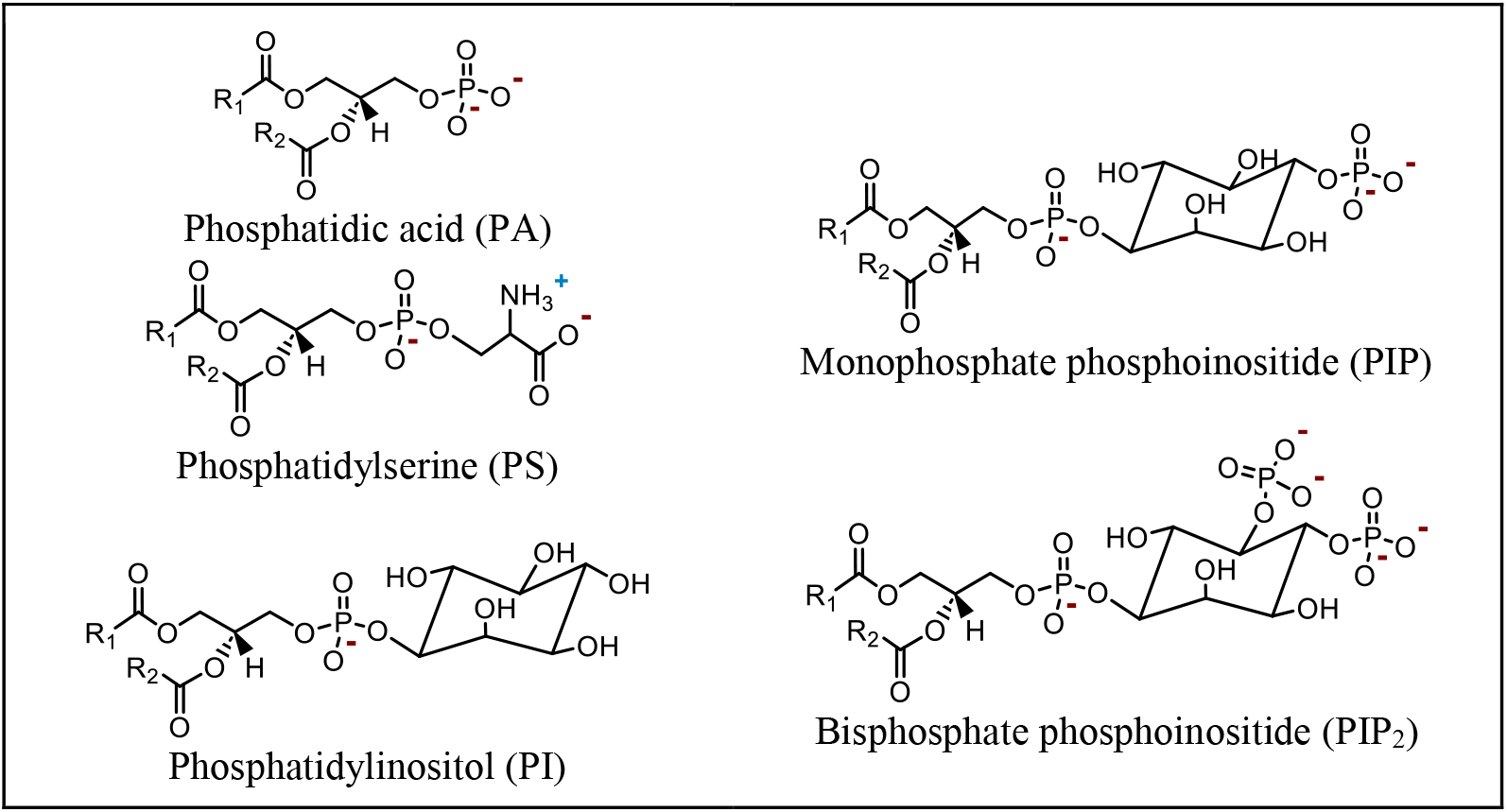
Structure of the anionic phospholipid classes. All anionic phospholipids are composed of a phosphate group attached to a glycerol backbone containing two esterified fatty acyl chains. Phosphatidic acid (PA) has the simplest structure. Phosphatidylserine (PS) and phosphatidylinositol (PI) refers to anionic phospholipids containing a serine or an inositol moiety linked to the phosphate group, respectively. PI that has one or two additional phosphate groups linked to the inositol are respectively names as monophosphate phosphoinositide (PIP) and bisphosphate phosphoinositide (PIP_2_)

The study of anionic phospholipids in plants is complicated because of their low abundance which requires the development of highly sensitive analytical methods. Indeed, PI, PA and PS respectively account for approximatively 7%, 2% and 1% of total phospholipids in *A. thaliana* leaves while the phosphoinositides account for less than 1% (Noack and Jaillais, 2020). Moreover, anionic phospholipids are categorized into structurally distinct classes (i.e. PA, PS, PI, PIP and PIP_2_) which all include a wide variety in fatty acyl chains both in length and saturation degree, leading to high differences in the hydrophobicity of the molecules that needs to be analyzed (Wang *et al*., 2016; Ogiso *et al*., 2008). A cost-efficient method was previously developed in which anionic phospholipids were separated by thin-layer chromatography (TLC) according to their characteristic head groups and, after separation, the associated fatty acids were analyzed using gas-chromatography (GC) (König *et al*., 2007; Heilmann and Heilmann, 2013). However, this method does not have the ability to separate individual molecular species as only the different classes of anionic phospholipids can be separated according to their polar head. Besides, it is not known where PS migrates in this method.

Importantly, little variations in the profile of anionic phospholipid species can lead to significant modulation of biological processes (Wang *et al*., 2016). Hence, there is an urgent need in the development of highly sensitive analytical methods for the comprehensive analysis of individual molecular species of each class of anionic phospholipids, notably from plant tissues. Ideally, this analytical method should be able to detect all anionic phospholipid species in one-shot. Mass spectrometry methods, by direct infusion or in combination with chromatography techniques, have emerged as reference methods for the analysis of phospholipids from plant samples (Lee *et al*., 2013; Wei *et al*., 2018; Cai *et al*., 2016; Wang *et al*., 2016). Among the different available chromatography methods, separation on reverse-phase columns by (ultra)-high performance liquid chromatography ((U)HPLC) are commonly performed. However, while the analysis of non-anionic phospholipids (PC, PE, …) with high sensitivity require little sample preparation and show high sensitivity and selectivity (Huang *et al*., 2019; Grison *et al*., 2015), difficulties remain as for the anionic phospholipids. Indeed, these molecules show low stability during MS analysis, which results in low analysis sensitivity, peak resolution problems, and therefore makes their analysis complicated (Lee *et al*., 2013; Wei *et al*., 2018). More specifically, the greatest peak tailing problems are encountered during the analysis of PA which is therefore very tricky to quantify in plant samples at physiological concentrations. Additionally, while phosphoinositide molecules can be analyzed by mass spectrometry methods without any derivatization, this comes with problems of recovery, stability and ion yields (Morioka *et al*., 2022). This makes their analysis very tricky when they are present at very low concentrations in plant samples. Different strategies have therefore been proposed to overcome these analytical problems, such as the addition of phosphoric acid to HPLC solvents, but these tips could not completely overcome peak resolution and sensitivity problems (Ogiso *et al*., 2008). More recently, different studies highlighted the high potential of anionic phospholipid derivatization and notably the methylation of hydroxyl groups, for their enhanced analysis by mass spectrometry (Cai *et al*., 2016; Canez *et al*., 2016; Wei *et al*., 2018; Zhang *et al*., 2019). By methylating the phosphate groups of anionic phospholipids, their negative charge would be neutralized (Wei *et al*., 2018; Wasslen *et al*., 2014), resulting in a significatively increased in analysis sensitivity and peak resolution (Huang *et al*., 2019; Wasslen *et al*., 2014; Wei *et al*., 2018; Lee *et al*., 2013). Different methylation strategies have been proposed, amongst which the use of trimethylsilyldiazomethane (TMSD) has now emerged (Yin *et al*., 2022) and was already used to comprehensively analyze PIP molecular species in plant membrane fractions (Ito *et al*., 2021). However, no method has been developed and applied yet for the simultaneous analysis and quantification of all anionic phospholipid species following their methylation from plant samples.

In this study, we aimed at filling this gap by developing a biphasic extraction with chemical modification to a targeted HPLC-MS/MS method for the simultaneous profiling of anionic phospholipids (PA, PS, PI, PIP and PIP_2_) from plant samples.

## 2. Results and discussion

### Optimization of HPLC and MS parameters for the profiling of methylated anionic phospholipids

Commercial mixtures of each investigated lipid class (PA, PS, PI, PIP and PIP_2_) were methylated and analyzed by direct infusion mass spectrometry for the confirmation of target adducts and fragments, as well as for the optimization of MS parameters (Declustering Potential DP, Collision Energy CE and Collision cell exit potential CXP) for their enhanced detection and fragmentation. It was decided to work in the positive ionization mode and ammonium formate was added to the solvents as it was shown to improve molecule ionization. Results in **Table 1** present the optimized MS ionization and fragmentation conditions. Besides, these results confirmed that the number of methyl groups added during the methylation reaction is specific to the polar head group and therefore to each anionic phospholipid class, as previously described (Lee *et al*., 2013). Indeed, while PIs were methylated once during the derivatization procedure, PAs and PSs were methylated twice; and PIPs and PIP_2_s were methylated 3 and 5 times, respectively. Our results also highlighted that the ionization occurs differently for the investigated anionic phospholipid classes as the main adducts were not the same for all molecules; the main detected adducts for PA and PI species were ammonium adducts; while protonated PS, PIP and PIP_2_ dominated the mass spectrum. Contrariwise, the fragmentation pattern was similar for each molecule as the primary detected fragments corresponded to the loss of the polar head group for all anionic phospholipids, which was in agreement with previous studies (Lee *et al*., 2013; Wasslen *et al*., 2014; Pi *et al*., 2016).

**Table 1.**
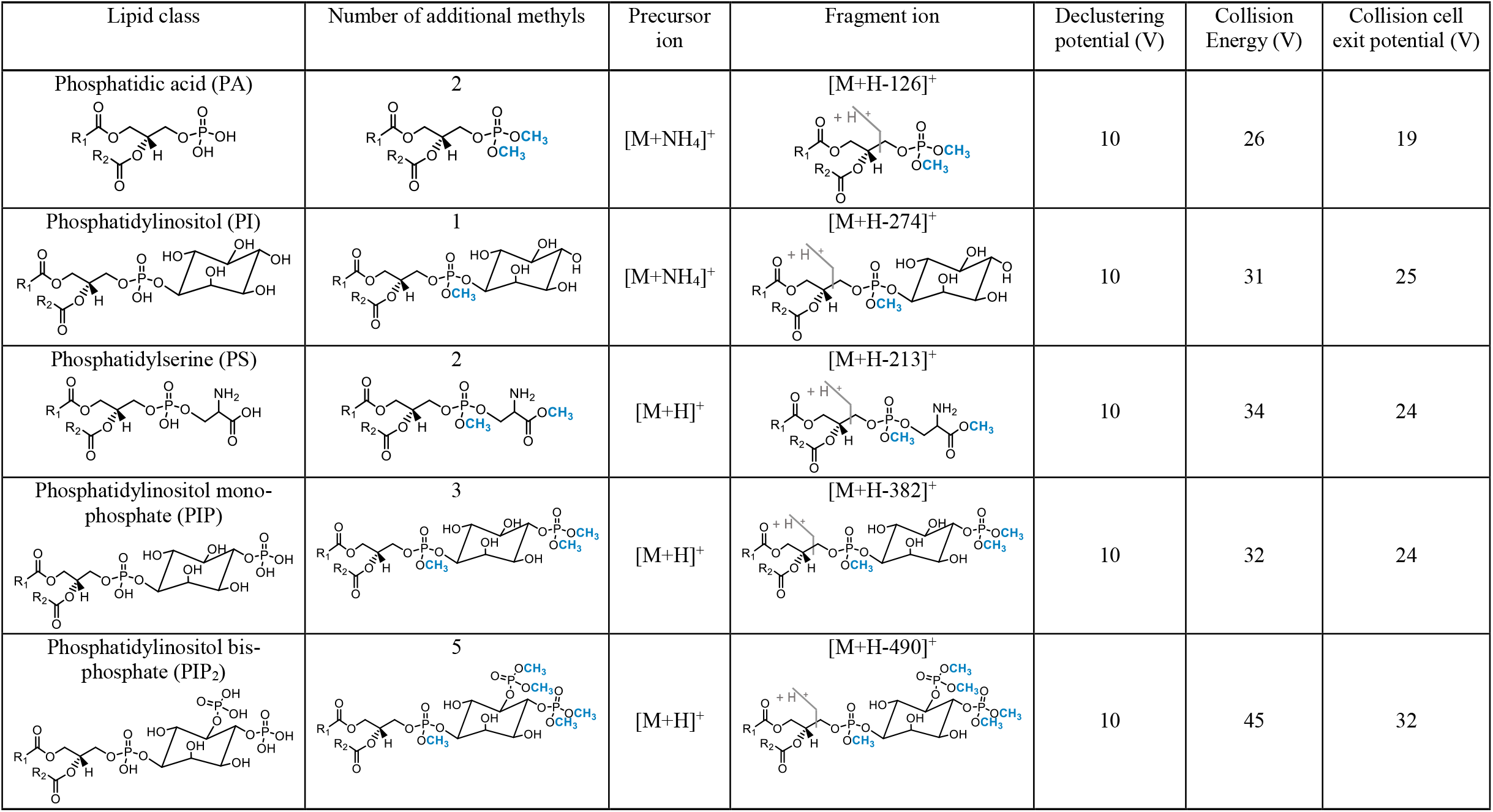
Optimized electrospray ionization - MS/MS analysis parameters for the analysis of methylated anionic phospholipids in the positive mode. For each anionic phospholipid class, the skeletal structure is presented before and after methylation reaction; as well as the main precursor ion, the fragmentation pattern, and the optimized conditions for their analysis by mass spectrometry.

| Lipid class | Number of additional methyls | Precursor ion | Fragment ion | Declustering potential (V) | Collision Energy (V) | Collision cell exit potential (V) |
| --- | --- | --- | --- | --- | --- | --- |
| Phosphatidic acid (PA) | 2 | [M+NH<sub>4</sub>]<sup>+</sup> | [M+H-126]<sup>+</sup> | 10 | 26 | 19 |
| Phosphatidylinositol (PI) | 1 | [M+NH<sub>4</sub>]<sup>+</sup> | [M+H-274]<sup>+</sup> | 10 | 31 | 25 |
| Phosphatidylserine (PS) | 2 | [M+H]<sup>+</sup> | [M+H-213]<sup>+</sup> | 10 | 34 | 24 |
| Phosphatidylinositol mono-phosphate (PIP) | 3 | [M+H]<sup>+</sup> | [M+H-382]<sup>+</sup> | 10 | 32 | 24 |
| Phosphatidylinositol bis-phosphate (PIP<sub>2</sub>) | 5 | [M+H]<sup>+</sup> | [M+H-490]<sup>+</sup> | 10 | 45 | 32 |

We then optimized the chromatographic resolution of the method using mixes of standard molecules for each of the investigated anionic phospholipid classes. These standards were first submitted to the methylation reaction, and the most appropriate analytical conditions were selected following analysis. We notably investigated the use of different column stationary phases (C4, C8 and C18 columns) and of different solvents, with or without the addition of ammonium formate that may enhance ionization of the molecules. Finally, we optimized the elution gradient with the selected solvents. The use of a C18 column, with a gradient of methanol:water (3:2) and isopropanol:methanol (4:1) allowed the best separation of the target methylated anionic phospholipids. Moreover, the addition of ammonium formate in solvents increased the analysis sensitivity and was therefore retained. Results presented in **Figure 2a** illustrate the chromatographic resolution of the method and notably the great peak shape for all investigated methylated anionic phospholipids. In comparison, poorer peak resolution, with mainly peak tailing and low sensitivity, was highlighted when non-methylated anionic phospholipids were analyzed using the same chromatographic parameters (**Figure 2b**). Notably, the peak asymmetry factor has been determined and shows that methylated PA, PI and PS species all have a great peak symmetry, with mean values of 0.99 for each of these three species when analyzed following methylation (**Figure 2c**). In comparison, the peak asymmetry factor of these same PA, PI and PS species when analyzed without being methylated beforehand confirmed the bad peak symmetry for PA (1.70) and PS (1.56) species, while the peak shape was good for PIs (0.95). Furthermore, the peak asymmetry factor of methylated PIPs (1.02) and PIP_2_s (1.16) also illustrate the greatest resolution of their peaks while these lipids could not be detected without the methylation step (**Figure 2c**).

**Figure 2.**
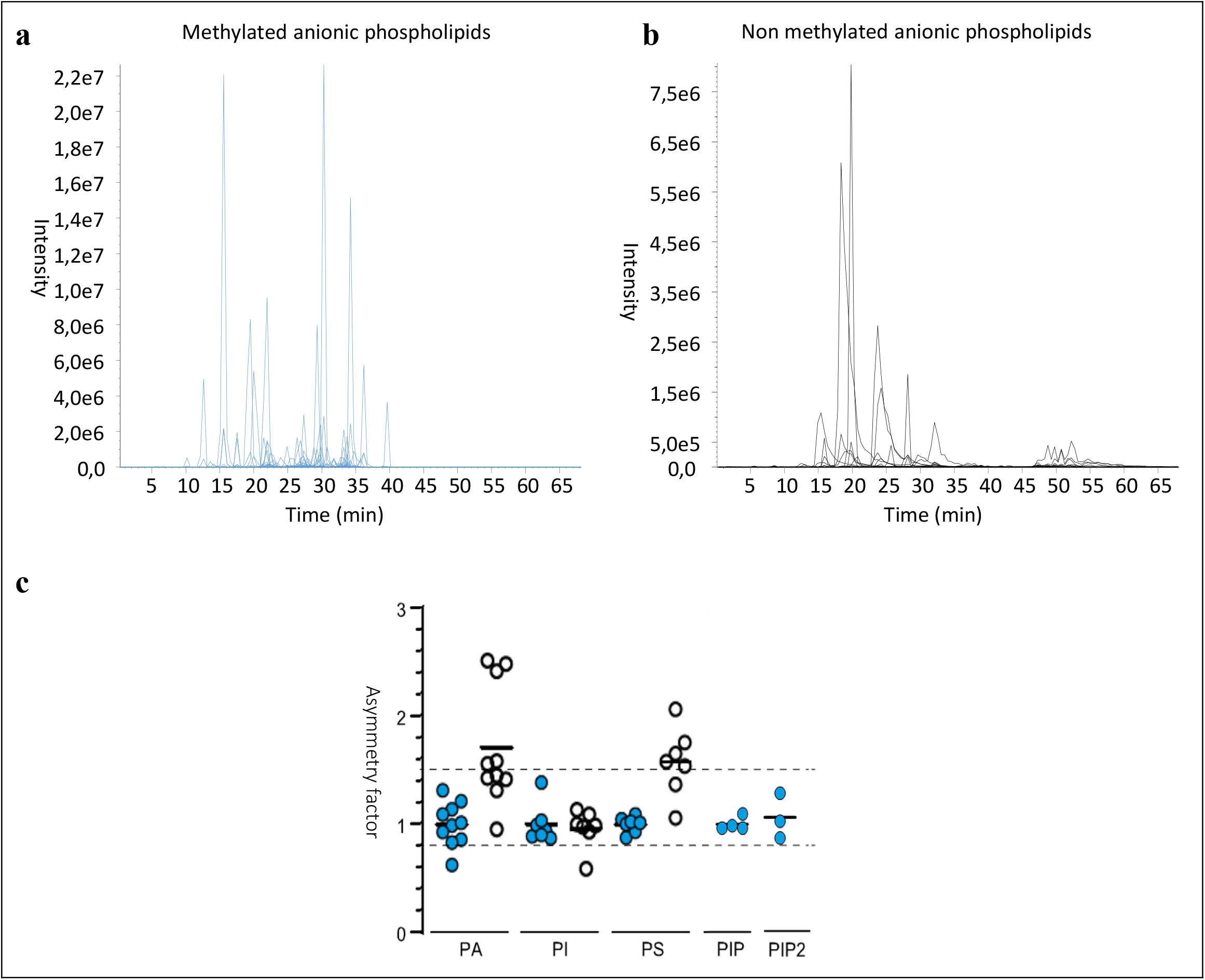
Presentation of the chromatographic resolution of the developed method. Samples containing mixtures of PA, PI, PS, PIP and PIP_2_ standards analyzed with the developed HPLC-MS/MS method following methylation (**a**) and without methylation (**b**). (**c**) For PA, PI and PS species, comparison of the peak asymmetry factor when samples were analyzed following (blue circles) or without (white circles) methylation. For PIP and PIP_2_, only the peak asymmetry factors are shown when samples were methylated, due to that no peaks were detected in absence of methylation.

As the ionization temperature is a determinant parameter when it comes to increase the sensitivity by mass spectrometry (Gao *et al*., 2016), we investigated the effect of the ionization temperature on the analysis of methylated anionic phospholipids. Mixes of standard molecules were thus analyzed by HPLC-MS/MS with the developed method and the optimized mass parameters, but with increasing ionization temperatures ranging from 200 to 600°C. Results presented in **Figure 3** showed that while methylated PAs and PIs requested lower temperatures for their ionization (respectively 200 and 250°C), higher temperatures allowed the optimal ionization of methylated PIPs and PIP_2_ (respectively 350 and 400°C), and PS species requested the highest ionization temperature (500°C). Our results therefore point out that the optimal ionization temperature highly differs between the investigated methylated lipid classes. In this view, choosing the optimal ionization temperature for all anionic phospholipid classes in the context of developing a single one-shot analysis for all these species by mass spectrometry is not possible. It was therefore chosen to select an ionization temperature of 350°C that was the best compromise to enhance sensitivity for all species and that was, in addition, close to the optimal ionization temperature of phosphoinositides species (PIP and PIP_2_). From all anionic phospholipids, the latter are present in the lowest quantities in plant samples (Noack and Jaillais, 2020) and therefore request the highest optimization of their analytical parameters so that they can be accurately detected and quantified.

**Figure 3.**
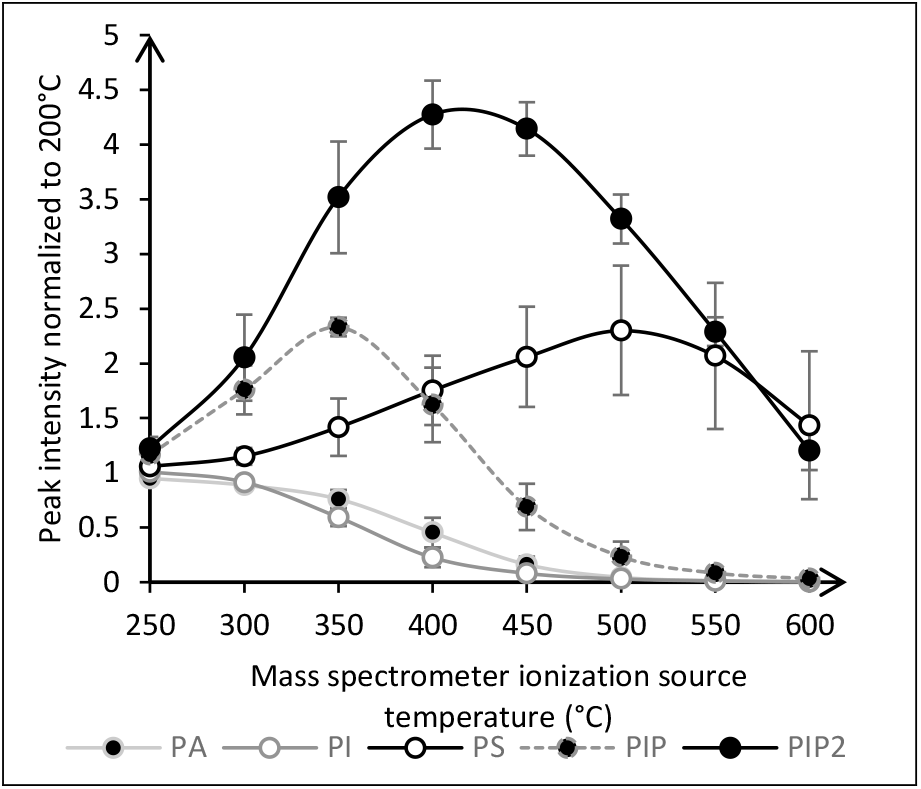
Optimization of the ionization temperature for the analysis of anionic phospholipids. Evaluation of the impact of the ionization temperature (200 to 600°C) on mass spectrometry signal intensity for PA, PI, PS, PIP and PIP_2_ lipids. The signal was normalized to that recorded at an ionization temperature of 200°C (n=3).

### Optimization of the methylation reaction conditions: evaluation of the impact of time and temperature

Different reaction conditions have been reported in the literature for the methylation of lipids using TMS-diazomethane, and notably the duration of the methylation and temperatures (Li *et al*., 2021). There was therefore a need in our method to optimize the reaction kinetics to select the best methylation conditions. In this context, we investigated the impact of the methylation time (ranging from 2 to 120 minutes) and of the reaction temperature (at 0, 23 and 40°C) on the methylation efficiency. The latter was determined by considering the percentage of the main methylated species on all reported species in the mixture following methylation.

First, the study of reaction temperature showed an increase in methylation efficiency between 0°C and 23°C, but low differences between 23°C and 40°C (**Figure 4a**). In light of these results, a temperature of 23°C was selected. In addition, results highlighted low differences in methylation efficiency in function of reaction time at the chosen temperature (**Figure 4b**), suggesting that the methylation reaction is almost instantaneous. Still, a slight decrease in methylation efficiency was noticed at 120 minutes of reaction for PI, PIP and PIP_2_ molecules, which was due to the increase in forms that were over-methylated. It is likely that methylation begins to occur on the inositol head groups of PIs, PIPs and PIP_2_ as this was not highlighted with PA and PS species. With regards to these results, we decided to select a reaction time of 10 minutes at 23°C. Besides, results also highlighted that, following methylation, the percentage of the main methylated form highly depends on the anionic phospholipid class that is methylated (**Figure 4c**). Indeed, while high differences were observed between the different investigated lipid classes, low differences were highlighted for lipids of a same class with different fatty acyl chains (**Table S1**).

**Figure 4.**
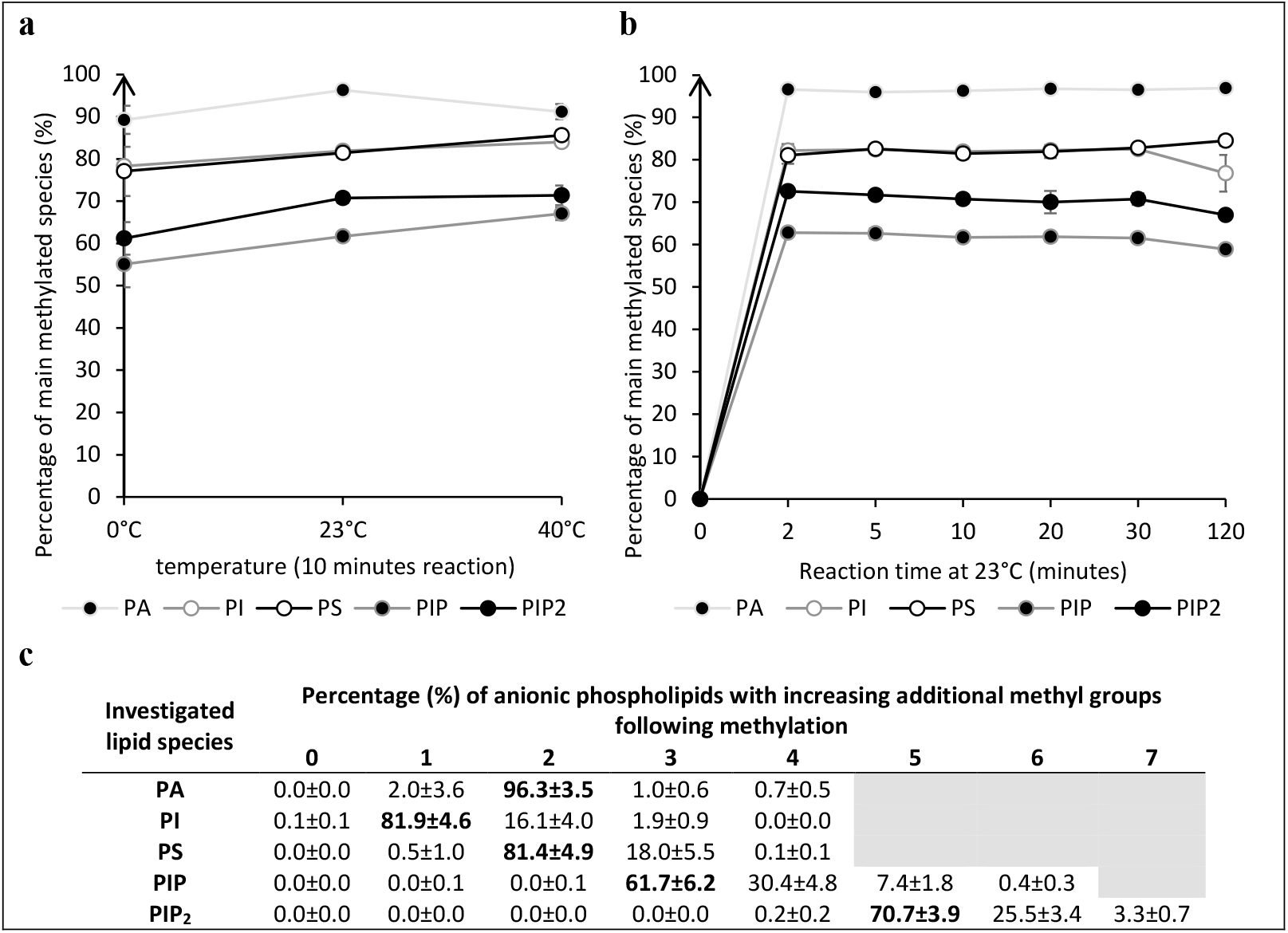
Optimization of the methylation reaction conditions. Impact of the methylation temperature at a constant reaction time of 10 min (**a**) and of the reaction time at a constant temperature of 23°C (**b**) on the percentage of the main methylated species for each analyzed class of anionic phospholipids (n=3). Monomethylated species were considered for PI; bismethylated species were considered for PA and PS. For the analysis of PIP and PIP_2_, species methylated 3 and 5 times were respectively considered. For a reaction time of 10 min at 23°C, the percentage of the investigated anionic lipid species methylated various times is presented (**c**; n=3). See **Table S1** for complete data.

### Analytical performance of the method

In order to confirm the high added value of the developed method and to evaluate its analytical performance, pure standards of anionic phospholipids were selected and analyzed at different concentrations with or without the methylation step. For these methylated and non-methylated species, the limit of detection (LOD) and limit of quantification (LOQ) were determined, as well as the linearity of the response and the peak width at 50% height.

Results presented in **Table 2** firstly demonstrate the high sensitivity of the developed method for all anionic phospholipid species investigated with LOD ranging from 0.27 to 11.82 fmol and LOQ ranging from 0.54 to 23.63 fmol, depending on the investigated lipids. The lowest LOD and LOQ values were obtained for the analysis of PA and the highest for PI(4,5)P_2_. With regard to the literature (Lee *et al*., 2013; Wei *et al*., 2018), these results show the improved sensitivity of our method as LODs of 7.5, 20 and 200 fmol were previously obtained for the analysis of PA, PS and PI using supercritical fluid chromatography coupled to a MS/MS analysis (Lee *et al*., 2013). Our method also demonstrates a highly increased sensitivity for the analysis of PA as compared to what has been previously reported by direct infusion MS (Wei *et al*., 2018). Moreover, our results confirmed that methylation of anionic phospholipids highly increases the sensitivity (**Table 2 and Figures 5, S1** and **S2**). The LOQ of PA was 128 times lower when the molecule was methylated in comparison with the non-methylated compound. For PS and PI species, the methylation allowed to decrease the LOQ by 4 and 32 times, respectively. PI4P and PI(4,5)P_2_ could not be detected without being methylated before analysis. PA and PS peak shape was also greatly improved by phospholipid methylation as the peak width at 50% height was decreased by more than 60% in comparison to not methylated samples. Additionally, the linear range was very wide for all anionic phospholipid species, with coefficients of determination (R^2^) higher than 0.99 for lipids that were methylated before analysis, and low coefficients of variation (Relative Standard Deviations RSDs).

**Table 2.** Analytical performance of the method. For each anionic phospholipid species analyzed during analytical method validation, the retention time, limits of detection (LOD) and of quantification (LOQ), linearity domain, coefficient of determination (R^2^) and peak width at 50% height are presented. Those data were presented for lipids that were methylated before analysis, as well as for species that were not methylated before analysis. The LOD and LOQ corresponded to the lowest concentration giving a signal/noise (S/N) higher than 3 and 10, respectively. “/” means no detectable signal. Calibration curves are presented in **Figures S1** and **S2**.

| Lipid species | Methylated or not | Retention time (min) | LOD (pg / fmol) | LOQ (pg / fmol) | LOQ fold decrease by methylation | Linearity domain (pg) | $R^2$ | Mean % RSD of calibration curve | width at 50% height |
| --- | --- | --- | --- | --- | --- | --- | --- | --- | --- |
| PA 18:0/18:1 | Methylated | 38.6 | 0.19 / 0.27 | 0.38 / 0.54 | 128 | 0.38 - 1562 | 0.9956 | 2.30 | 0.34 |
|  | Not methylated | 54.0 | 12.21 / 17.37 | 48.83 / 69.46 |  | 48.83 - 12500 | 0.9974 | 8.22 | 2.73 |
| PS 16:0/18:1 | Methylated | 24.4 | 3.05 / 4.00 | 12.21 / 16.02 | 4 | 12.21 - 3125 | 0.9958 | 2.46 | 0.53 |
|  | Not methylated | 26.4 | 24.41 / 32.03 | 48.83 / 64.08 |  | 48.83 - 3125 | 0.9895 | 6.26 | 1.40 |
| PS 18:0/18:2 | Methylated | 27.0 | 3.05 / 3.87 | 12.21 / 15.49 | 4 | 12.21 - 3125 | 0.9979 | 2.45 | 0.48 |
|  | Not methylated | 28.9 | 24.41 / 30.97 | 48.83 / 61.96 |  | 48.83 - 3125 | 0.9888 | 6.49 | 1.25 |
| PI 16:0/18:1 | Methylated | 26.1 | 3.05 / 3.64 | 6.10 / 7.29 | 32 | 6.10 - 3125 | 0.9913 | 5.25 | 0.51 |
|  | Not methylated | 23.7 | 97.66 / 116.67 | 195.31 / 233.32 |  | 195.31 - 12500 | 0.9882 | 11.54 | 0.61 |
| PI 17:0/14:1 | Methylated | 15.4 | 3.05 / 3.84 | 6.10 / 7.67 | 32 | 6.10 - 3125 | 0.9973 | 5.19 | 0.40 |
|  | Not methylated | 13.6 | 97.66 / 122.84 | 195.31 / 245.67 |  | 195.31 - 12500 | 0.9970 | 10.64 | 0.46 |
| PI4P 17:0/20:4 | Methylated | 25.0 | 3.05 / 3.20 | 6.10 / 6.40 | / | 6.10 - 3125 | 0.9906 | 4.52 | 0.72 |
|  | Not methylated | / | / | / |  | / | / | / | / |
| PI(4,5)P<sub>2</sub> 17:0/20:4 | Methylated | 24.7 | 12.21 / 11.82 | 24.41 / 23.63 | / | 24.41 - 3125 | 0.9878 | 3.75 | 0.49 |
|  | Not methylated | / | / | / |  | / | / | / | / |

**Figure 5.**
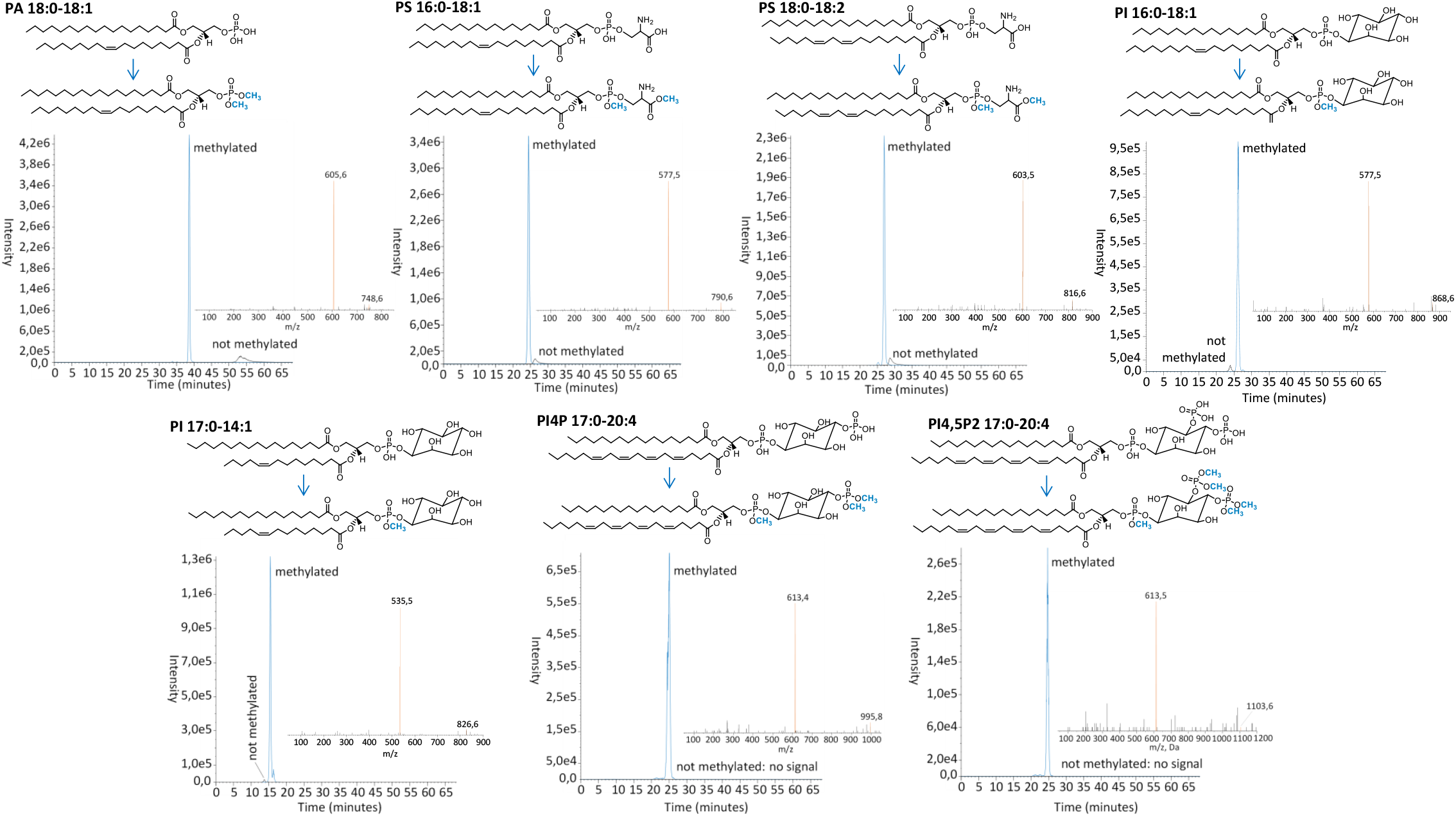
Presentation of the main species following methylation of each anionic phospholipid analyzed during analytical method validation. Chromatograms show comparisons of the HPLC-MS signal before and after derivatization. The peak in blue corresponds to the signal of the main species analyzed following sample methylation while the signal in black corresponds to the unmethylated species of unmethylated samples at the same concentration. Of note, signals were below the limit of detection for unmethylated PI4P and PI(4,5)P_2_ lipids. Mass spectra show fragments of the target adducts for each molecule. The main fragment targeted for MS/MS analysis was highlighted in orange.

### Application of the method to plant samples

To validate our method, we performed the extraction and the quantification of anionic phospholipids on 7 different biological materials from 3 different plant species that were either monocots or dicots. For the dicot plants, we chose *Arabidopsis thaliana* in which we sampled 4-weeks old leaves, 7-days old seedlings either took as a whole or split into aerial organs (cotyledons and primary leaves) from one side and root organs on the other side, as well as Landsberg *erecta* suspension cells 10-days old after sub-culturing. We also sampled 4-weeks old leaves of the dicot *Nicotiana benthamiana*. For the monocot plants, we choose to sample 1-week old leaves from *Zea mays*. Approximately 30 mg of fresh weight plants were used for each analysis.

Interestingly, results presented in **Figures 6, S3** and **S4** firstly illustrate the low variability of the results regarding the analysis of different biological replicates. Indeed, when looking at the main species of each anionic phospholipid class analyzed from mature leaves of *Arabidopsis thaliana*, we observed a very low mean RSDs of 3.92, 4.33, 4.14 and 7.17 % for PA (34:2, 34:3, 36:4 and 36:5), PI (34:2 and 34:3), PIP (34:2, 34,3) and PIP_2_ (34:2, 34:3) species, respectively. Slightly higher RSD were observed for the analysis of PS species (36:4, 36:5) with a mean value of 18.60%.

**Figure 6.**
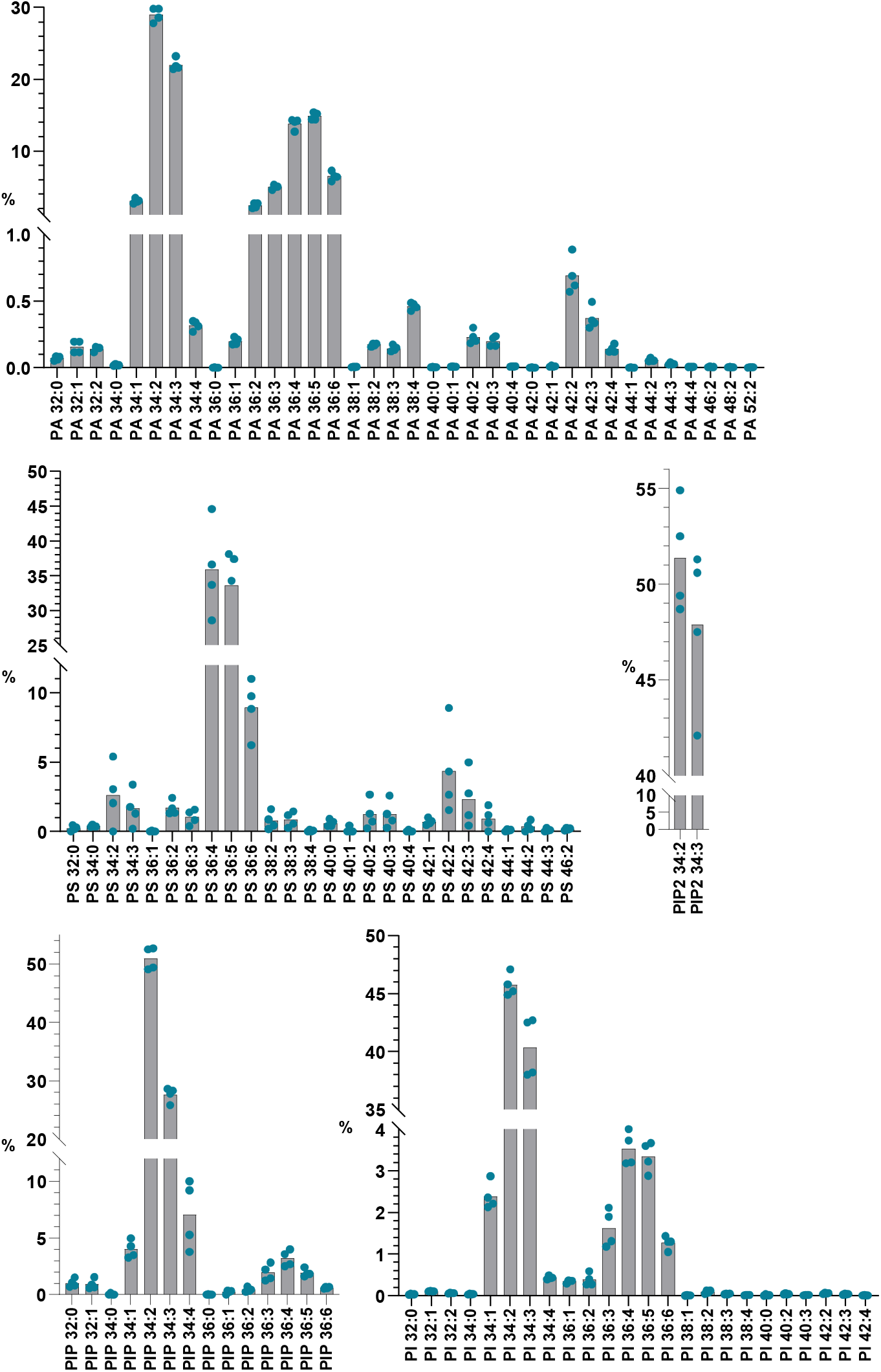
Composition of anionic phospholipids in *Arabidopsis thaliana* leaves. Data are expressed in % of total detected species. n = 4 biological replicates, the dots show the dispersion of data. See **Figures S3** and **S4** for data relative to other plant species and organs.

When comparing the anionic lipids extracted from mature leaves of *A. thaliana, N. benthamiana* and *Z. mays* (**Figure 7**), we observed a very similar PA composition with PA 34:2 and 34:3 being the primary major molecular species and PA 36:4 and 36:5 secondary major species. Similarly, PI has PI 34:2 and 34:3 as primary major species, but does not have secondary major species as for PA (**Figure 7**). These results are similar to what was published previously for the fatty acid content of PA and PI in *A. thaliana* (Devaiah *et al*., 2006). Our results further show that the phosphoinositide family consisted of monophosphate phosphoinositides PIP 34:2 and 34:3 in all 3 leaf samples, with the notable presence of PIP 32:0 and 34:4 in *Z. mays* leaves, and bis-phosphate phosphoinositides PIP_2_ 34:2 and 34:3 in all samples (**Figure 7**). These results are similar to what was previously published for the phosphoinositide pool extracted from the Arabidopsis rosette leaves and analyzed by TLC coupled to GC (Devaiah *et al*., 2006; König *et al*., 2007). The PIP and PIP_2_ molecular species are the same than the ones detected in the pool of PI, which indicates that PIP and PIP_2_ are derived from PI and are not further edited, except for the PIP of the monocot *Z. mays* where some acyl-chain editing could occur according to the profiles obtained in this study. In contrast to PA, PI and phosphoinositides, the PS molecular species are much more diverse in the mature leaves of *A. thaliana, N. benthamiana* and *Z. mays* (**Figure 7**). In the *A. thaliana* mature leaves, the major species were PS 36:4 and 36:5 while several additional species were also detected in *N. benthamiana* such as PS 38:3, 36:3 and 34:3. Interestingly and in contrast to the dicots species, the monocot *Z. mays* clearly displays very long chain fatty acids (VLCFA) in the PS pool up to PS 42:2 and 42:3.

**Figure 7.**
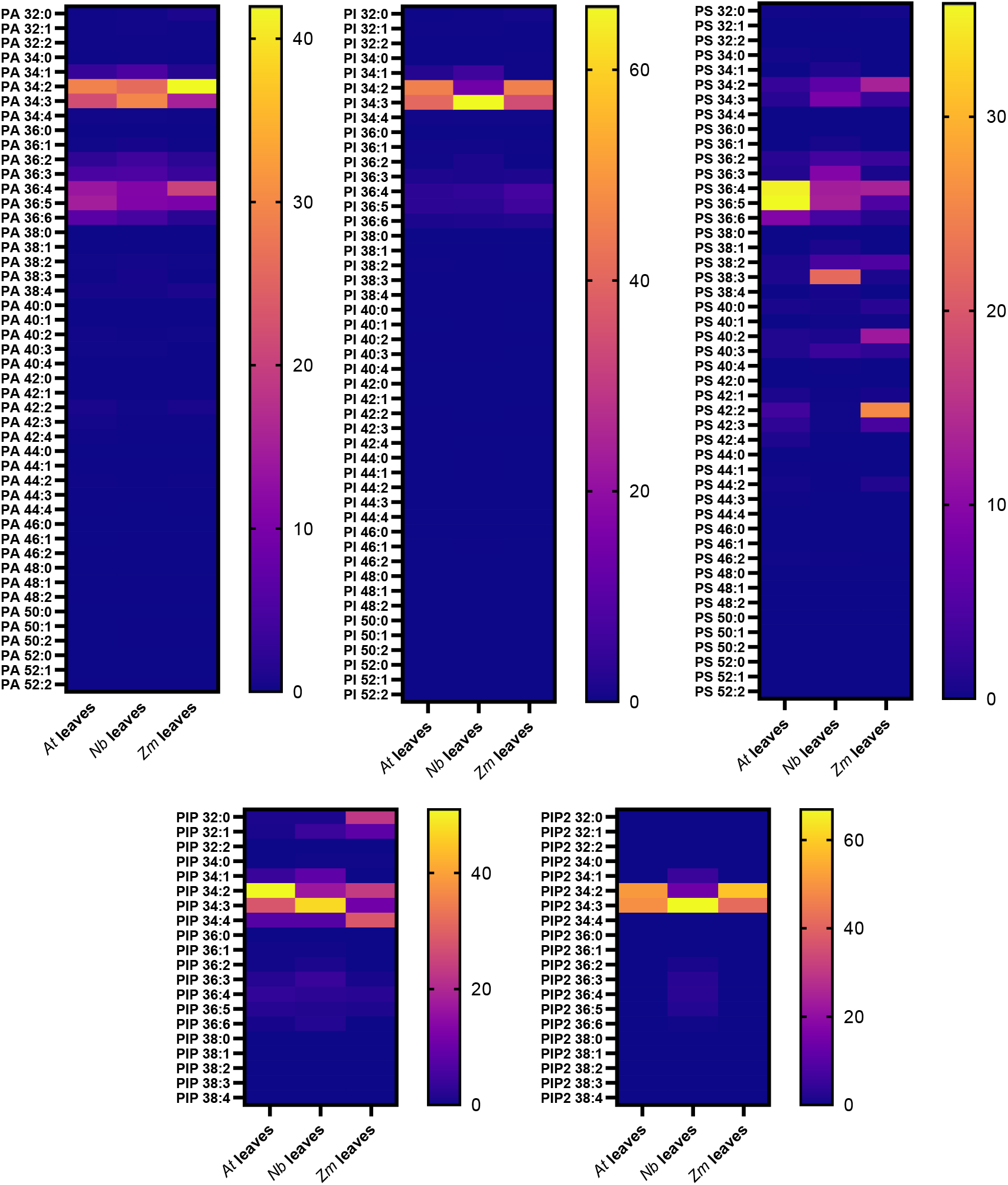
Heatmap presenting the proportions in anionic phospholipids analyzed with the developed method in leaves from different plant species. *At: Arabidopsis thaliana*; *Nb: Nicotiana benthamiana*; *Zm: Zea mays*. Data are expressed in % and are means from 4-5 measurements.

However, the molecular species of anionic phospholipids found in mature leaves might not be representative of the whole plant, and might, in other words differ from one organ to another. We therefore analyzed the composition of the anionic lipids found in different organs of *A. thaliana*. For that purpose, we chose to analyze the composition of the anionic lipids in either the aerial part (cotyledons and primary leaves) or the roots of 7-day old dissected seedlings. As a reference, we also analyzed 7-days old whole seedlings that were not dissected and, for the comparison, we put the results previously obtained in **Figure 7** (mature leaves). We additionally analyzed the content of undifferentiated *Arabidopsis* suspension cell culture as this would be a great comparison between cells embedded within a differentiated organ and cells immortalized that grow in a liquid media. However, it should be noted that the analysis of the different plant organs was made in the Colombia-0 (Col-0) ecotype while the cell culture is coming from the Landsberg *erecta* (L*er*) ecotype. Thus, the differences observed might also be due to the ecotype background. Interestingly, our results show that PA, PI, PIP and PIP_2_ composition was basically similar to what we observed in *A. thaliana* leaves in Figure 7 (**Figure 8**). However, the PS content strikingly differs among the different *A. thaliana* organs (**Figure 8**). Indeed, the roots were particularly enriched in VLCFA-containing PS (42:2, 42:3 and to a lesser extend 40:2 and 40:3) as compared to the aerial organs. In whole seedlings, VLCFA-containing PS were not easily visible. This is probably due to the dilution of membranes coming from the roots in the total pool of membranes coming from the aerial organs, as the latter are much more abundant than in the roots. It is interesting to note that the cell culture also displays a VLCFA-containing PS profile, similar to the root samples. Our results are consistent with other previously published studies on roots or whole seedlings (Devaiah et al., 2006; Platre et al., 2018).

**Figure 8.**
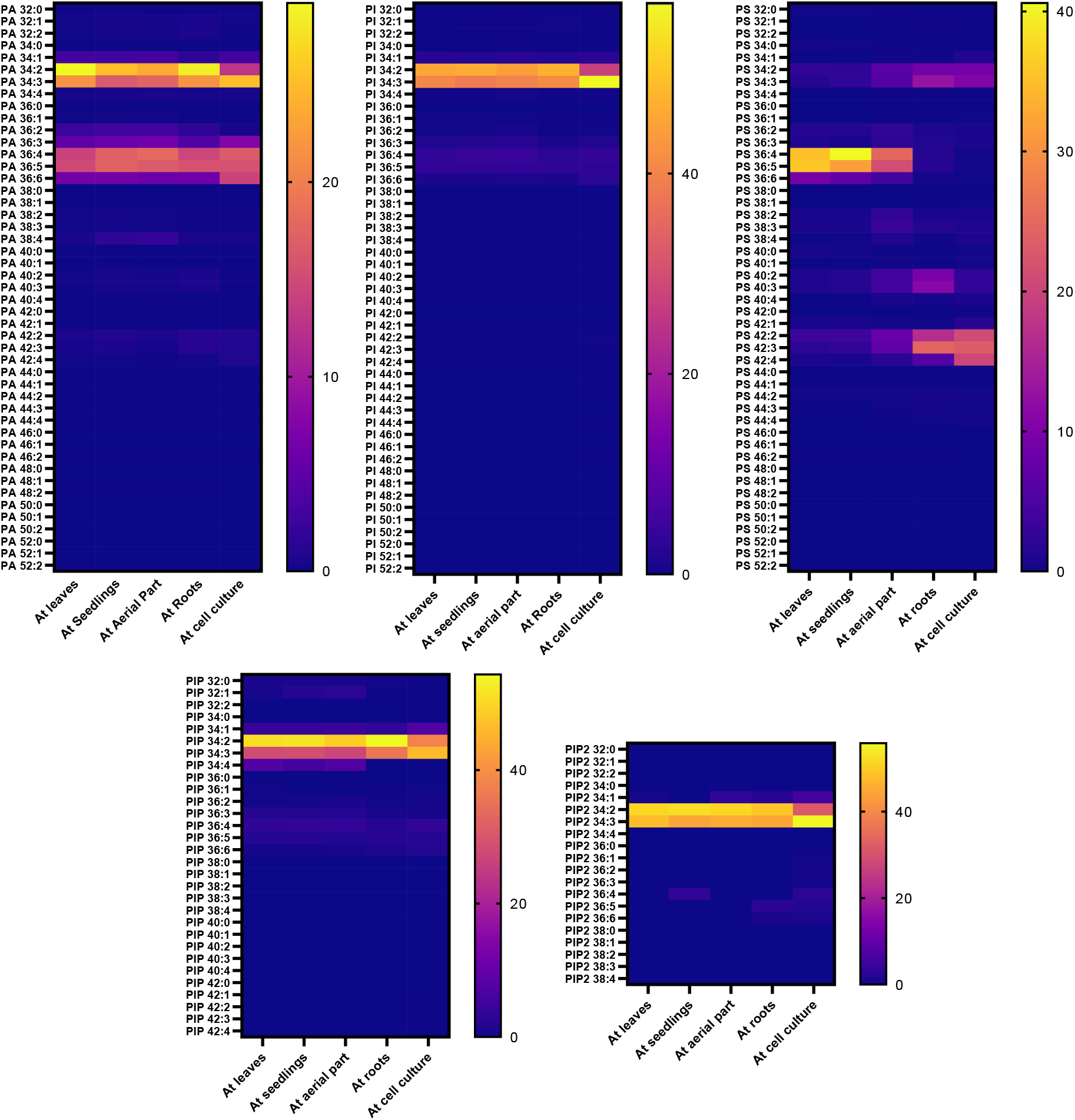
Heatmap presenting the proportions in anionic phospholipids analyzed with the developed method in different organs of *Arabidopsis thaliana. At*: *Arabidopsis thaliana*. Aerial part corresponds to cotyledons and primary leaves. Data are expressed in % and are means from 4-5 measurements.

### General discussion

The poor chromatographic resolution of anionic phospholipids, as well as their low abundance in biological matrices and weak stability during MS analysis, are the main factors making the analysis of these lipids complicated (Noack and Jaillais, 2020). Furthermore, their distribution into structurally different classes (i.e. PA, PS, PI, PIP and PIP_2_) with a high diversity in fatty acyl chain length and saturation degree inside each class makes the analysis even more complex. Previous studies already pointed out the high potential of lipid methylation, notably using TMS-diazomethane, as a derivatization procedure that increases the molecule stability during its ionization and fragmentation (Wasslen *et al*., 2014; Lee *et al*., 2013; Clark *et al*., 2011), therefore increasing the sensitivity of their analysis by mass spectrometry. However, to the best of our knowledge, the present method is the first developed analytical method for the profiling and accurate quantification of all anionic phospholipids classes from plant tissues in one-shot.

In our approach, we indeed propose an effective strategy for the methylation of all anionic phospholipid species from plant samples using TMS-diazomethane, as well as a highly sensitive method for their analysis by HPLC-MS/MS. This was achieved by optimizing the methylation reaction conditions, HPLC analytical separation and MS ionization and fragmentation parameters. Our results showed a great improvement of analysis sensitivity and resolution, with an improved peak shape, in comparison with the analysis of the same lipids without any methylation step during the sample preparation procedure. In particular, methylation largely enhanced the sensitivity of the analysis of PA and PS species, which presented very large peaks when being analyzed without methylation. Large peaks can hardly be quantified due to their bad peak shapes and therefore be a source of high variation in the obtained signal. Our method now solves this issue and allows the detection of PA and PS with a high accuracy. Given the low abundance of anionic phospholipids, this parameter is crucial in the confidence we grant to the results. The increase in the accuracy or the confidence threshold with our method is even more important for the phosphoinositides PIP and PIP_2_ as their abundance is even lower than PA or PS (Noack and Jaillais, 2020). We also confirmed the applicability of our method to the analysis of plant biological samples with a low variability for all analyzed anionic phospholipid classes and for all plant matrices investigated, including leaves of three different plant species (*A. thaliana, N. benthamiana* and *Z. mays*) and different *A. thaliana* organs.

The anionic phospholipids require a very precise, sensitive and accurate method of quantification as they are highly effective at very low quantities in membranes and accumulate at define spots of the endomembrane system in a transient manner where they are quickly metabolized in other lipid forms. Here we provide a HPLC-MS/MS method that not only confirmed previous published results on the composition of individual classes of anionic phospholipids, but also substantially improved their detection and quantification altogether in one-shot. Additionally, our results provide new evidences towards the differential fatty acid composition of PS in roots versus aerial organs. The presence of VLFCA in PS seems quite specific to the plant kingdom as already described in several publications using TLC-GC (Yamaoka *et al*., 2011; Yamaoka *et al*., 2021) or LC-MS/MS without derivatization of lipids (Devaiah *et al*., 2006; Maneta-Peyret *et al*., 2014). Of note, animal PS mostly contains 18:0-20:4 fatty acids (Lorent *et al*., 2020). The results on our plant samples suggest that PS is regulated during its synthesis or is subjected to acyl-chain editing after its synthesis to fulfill different functions according to the physiological need of a given organ. Indeed, it has been shown previously that the subcellular localization of PS at either the plasma membrane or endosomes varies during root cell differentiation and that this gradient of localization tunes auxin signaling through the nano-clustering of the small GTPase ROP6 (Platre *et al*., 2019). Therefore, measuring the level of anionic lipids, including PS, and their fatty acid content with a high level of precision and accuracy during cell differentiation or in different membrane compartments of the cell is important and may be a leverage to understand the regulation of signaling during development. Lastly, the development of our method opens the way to the implementation of analytical strategies for the quantification of other lipids classes in plant samples, and notably for the analysis of lyso forms of anionic phospholipids, which are phospholipids containing one single fatty acyl chain. As these lipids are known to accumulate under a wide range of biological conditions, and notably in response to biotic and abiotic stresses (Hou *et al*., 2016; Okazaki and Saito, 2014), the improvement of analytical methods for their more accurate and sensitive quantification would assuredly contribute to a better understanding of their roles in plant cell processes.

## 3. Experimental procedures

### Chemicals and Reagents

HPLC-grade methanol, chloroform, isopropanol and formic were purchased from Sigma-Aldrich. TMS-diazomethane (2M in hexane) was purchased from Fischer. The following lipids used for the experiments were all purchased from Avanti Polar Lipids: egg PA, soybean PS, brain PI4P, brain PI(4,5)P_2_, soybean PI. The following synthetic standard molecules were also bought from Avanti Polar Lipids: PA 18:0/18:1 and 15:0/18:1, PI 17:0/14:1 and 16:0/18:1, PS 18:0/18:2, 16:0/18:1 and 17:0/17:0, PI4P 17:0/20:4 and PI(4,5)P_2_ 17:0/20:4. All lipids were stored at -20°C before sample preparation. The internal standards were added in the samples at a final concentration of 0.1 µg/mL.

### Lipid extraction and derivatization procedure

725 µL of a MeOH:CHCl_3_:1M HCl (2:1:0.1 ; by volume) solution and 150 µL water were added to the samples in Eppendorf tubes which were then grinded three times for 30 seconds using a TissueLyser II (Qiagen^®^) after addition of metal beads. Following the addition of the internal standard when appropriate and of 750 µL CHCl_3_ and 170 µL HCl 2M, the samples were vortexed and centrifuged (1500 g / 5 min). The lower phase was washed with 708 µL of the upper phase of a mix of MeOH:CHCl_3_:0.01M HCl (1:2:0.75; by volume). Samples were vortexed and centrifuged (1500 g / 3 minutes) and the washing was repeated once. Then, samples were kept overnight at -20°C.

The organic phase was transferred to a new Eppendorf tube and the methylation reaction was carried out with 50 µL of TMS-diazomethane (2M in hexane). After 10 minutes of reaction at 23°C, the reaction was stopped by adding 6 µL of glacial acetic acid. 700 µL of the upper phase of a mix of MeOH:CHCl_3_:H_2_O (1:2:0.75; by volume) was added to each sample which was then vortexed and centrifuged (1500 g/ 3 minutes). The upper phase was removed and the washing step was repeated once. Finally, the lower organic phases were transferred to a new Eppendorf tube. Following the addition of 100 µL MeOH:H_2_O (9:1; by volume), the samples were concentrated under a gentle flow of air until only a drop remained. 80 µL of methanol were then added to the samples, which were submitted to ultrasounds for 1 minute, before adding 20 µL water, and were submitted to 1 more minute sonication. The samples were finally transferred to HPLC vials for analysis.

### HPLC-MS/MS analysis

Analysis of methylated and unmethylated anionic phospholipids was performed using a high performance liquid chromatography system (1290 Infinity II, Agilent) coupled to a QTRAP 6500 mass spectrometer (Sciex). The chromatographic separation of anionic phospholipids was performed on a reverse phase C18 column (SUPELCOSIL™ ABZ PLUS; 10 cm x 2.1 mm, 3µm, Merck) using methanol:water (3:2) as solvent A and isopropanol:methanol (4:1) as solvent B at a flow rate of 0.2 mL/min. All solvents were supplemented with 0.1% formic acid and 10 mM ammonium formate. 10 µL of samples were injected and the percentage of solvent B during the gradient elution was the following: 0–20 min, 45%; 40 min, 60%; 50 min, 80%; 52 min, 100%; 61 min, 100%; 61.1-68 min, 45%. The column temperature was kept at 40°C. Nitrogen was used for the curtain gas (set to 35), gas 1 (set to 40), and gas 2 (set to 40). Needle voltage was at +5500 V with needle heating at 350°C; the declustering potential (DP) was +10 V. The collision gas was also nitrogen; collision energy (CE) was set between 26 to 45 eV according to the lipid classes (**Table 1**). The dwell time was set to 5 ms. Mass spectrometry analysis was performed in the positive ionization mode.

### Method development

The optimization of mass spectrometer parameters for the analysis of anionic phospholipids, and more specifically of the DP, CE and collision cell exit potential (CXP), was performed by direct infusion mass spectrometry using methylated mixes of standards (egg PA, soy PS, brain PI4P, brain PI(4,5)P_2_, soy PI). For each sample, 0.1 mg of lipid was methylated as described above and resuspended in 1 mL of methanol:H_2_O 8:2 containing 0.2 % of formic acid and 6.5 mM ammonium formate for ionization, before being directly infused at a flow rate of 5 µL/min in to the QTRAP 6500 mass spectrometer (Sciex).

The optimization of the chromatographic analysis of anionic phospholipid was performed using a mixture containing 0.01 mg of each mix of standard molecule (egg PA, soybean PS, brain PI4P, brain PI(4,5)P_2_, soybean PI). The mixture was methylated as described above and injected by HPLC-MS/MS using different HPLC columns, solvents and solvent gradients. The chromatographic conditions allowing the most optimal chromatographic resolution for targeted anionic phospholipids were selected, as well as the optimized mass spectrometer analysis parameters. Investigated anionic phospholipid species were the following: PA (32:0, 32:1, 32:2, 34:0, 34:1, 34:2, 36:1, 36:2, 38:4, 38:5, 38:6, 40:6), PI (34:1, 34:2, 34:3, 36:1, 36:2, 36:3, 36:4, 36:5), PS (34:1, 34:2, 34:3, 36:1, 36:2, 36:3, 36:4, 36:5, 36:6), PIP (32:0, 34:0, 34:1, 36:1, 36:2, 36:3, 36:4, 38:2, 38:3, 38:4, 38:5, 40:4) and PIP_2_ (32:0, 34:0, 34:1, 36:1, 36:2, 36:3, 36:4, 38:3, 38:4, 38:5, 40:4, 40:5). See **Table S2** for the list of MRM information (Q1 and Q3 mass, DP, CE and CXP).

The optimization of ionization temperature for the analysis of anionic phospholipids was performed using the same mixture containing 0.01 mg of each mix of standard molecule methylated as described above. These were analyzed with the selected optimal chromatographic conditions but with increasing ionization temperatures (200 – 250 – 300 – 350 – 400 – 450 – 500 – 550 – 600 °C). All analyses were repeated three times. The most optimal ionization temperature was determined for each anionic phospholipid class. See **Table S2** for the list of MRM information (Q1 and Q3 mass, DP, CE and CXP).

The impact of time and temperature on the methylation reaction has been studied by performing the methylation reaction on the same mixture of standards as detailed before, but by investigating different methylation temperatures (0 – 23 and 40°C) and reaction duration (0 – 2 – 5 – 10 – 20 – 30 – 120 minutes). All analyses were repeated three times. See **Table S3** for the list of MRM information (Q1 and Q3 mass, DP, CE and CXP).

### Method validation

For the validation of the developed method, synthetic standard molecules were used (18:0/18:1 PA, 17:0/14:1 PI, 16:0/18:1 PI, 18:0/18:2 PS, 16:0/18:1 PS, 17:0/20:4 PI4P and 17:0/20:4 PI(4,5)P_2_). Methylated 17:0/17:0 PS was used as the internal standard. See **Table S4** for the list of MRM information (Q1 and Q3 mass, DP, CE and CXP).

The limit of detection (LOD), limit of quantification (LOQ) and linearity domain of each of these anionic phospholipids were determined following methylation with the optimized methylation conditions; as well as without any methylation. Methylated 17:0/17:0 PS was used as an internal standard at a concentration of 0,1 µg/mL in all of these samples. LOD and LOQ corresponded to the injected quantity giving a signal-to-noise ratio (S/N) of 3 and 10, respectively. For the assessment of the linearity domain, increasing quantities of methylated and non-methylated synthetic standard molecules were prepared and analyzed by HPLC-MS/MS with the developed method. The peak asymmetry factor was determined by dividing, at 10% peak height, the peak duration from the peak top to its end point by the peak duration from the peak starting point until the peak top, as previously described (Kantz *et al*., 2019). All analyses were repeated three times.

### Application of the method to biological material

The extraction and semi-quantification of anionic phospholipids was carried out using the developed method to the analysis of different biological materials: (i) leaves from 4-week old *A. thaliana*; (ii) leaves from 4-week old *N. benthamiana*; (iii) leaves from 1-week old *Z. mays* cultivated on soil; (iv) entire seedlings, (v) aerial part and (vi) roots from *A. thaliana* seedlings cultivated *in vitro* for 7 days on MS plates; (vii) *A. thaliana* var. *Landsberg erecta* suspension cells collected 10 days following subculture. Approximately 30 mg of fresh mass was used for each analysis and 15:0/18:1 PA was used as the internal standard. All analyses were repeated 4-5 times. See **Table S5** for the list of MRM information (Q1 and Q3 mass, DP, CE and CXP).

### Data analysis

HPLC-MS data were treated using the Analyst (AB Sciex 1.6.2) and MultiQuant (AB Sciex 3.0.3) softwares.

## 4. Acknowledgments

We thank Bordeaux-Metabolome platform for lipid analysis (https://www.biomemb.cnrs.fr/en/lipidomic-plateform/) supported by Bordeaux Metabolome Facility-MetaboHUB (grant no. ANR–11–INBS–0010 to SM and LF).

Manon Genva thanks the National Fund for Scientific Research (FNRS, Belgium, through grant 2022/V 3/5/053-40010130-JG/JN-2724) and the University of Liège for mobility grants. This work was supported by the French National Research Agency (project ANR PRC PlayMobil, grant no. ANR-19-CE20-0016 to YB, LF and SM, LF and project ANR PRC CALIPSO grant no. ANR-18-CE13-0025 to YB and LF). Louise Fougère thanks the MESRI fellowship for her PhD. We thank Louise Vilain for manuscript proof-reading.

## 5. Short legends for Supporting Information

**Figure S1**. Presentation of the calibration curves for methylated anionic phospholipids species. Each point corresponds to the mean of 3 repetitions.

**Figure S2**. Presentation of the calibration curves for non-methylated anionic phospholipids species. Each point corresponds to the mean of 3 repetitions. Of note, no signal was detected for non-methylated PIP and PIP_2_ lipids.

**Figure S3**. Composition of anionic phospholipids in *Nicotiana benthamiana* leaves **(a)** and *Zea mays* leaves **(b)**. Data are expressed in %, n = 5 biological replicates, the dots show the dispersion of data.

**Figure S4**. Composition of *Arabidopsis thaliana* anionic phospholipids in 7-days old seedlings **(a)**, aerial part (cotyledons and primary leaves) **(b)**, roots **(c)** and suspension cells 10-days after sub-culturing **(d)**. Data are expressed in %, n = 5 biological replicates, the dots show the dispersion of data.

**Table S1**. For a reaction time of 10 min at 23°C, presentation of the percentage of the investigated anionic lipid species methylated various times (n=3).

**Table S2**. List of MRM information (Q1 and Q3 mass, DP, CE and CXP) used for the analysis of anionic PL commercial mixes in the context of the optimization of chromatographic separation and of ionization temperature.

**Table S3**. List of MRM information (Q1 and Q3 mass, DP, CE and CXP) used for the analysis of anionic PL commercial mixes in the context of the optimization of the methylation reaction conditions.

**Table S4**. List of MRM information (Q1 and Q3 mass, DP, CE and CXP) mass used for the confirmation of the analytical performance of the method

**Table S5**. List of MRM information (Q1 and Q3 mass, DP, CE and CXP) used for the analysis of plant samples

## 7. Author contribution statement

L. Fouillen conceptualize the study and the analytical flowchart. All authors performed the extraction, derivatization methods on lipid standards and biological samples. MG and L. Fouillen performed the LC-MS^2^ analysis. All authors wrote and corrected the manuscript.

