## Supporting information for "A global LC-MS^2^-based methodology to identify and quantify anionic phospholipids in plant samples"

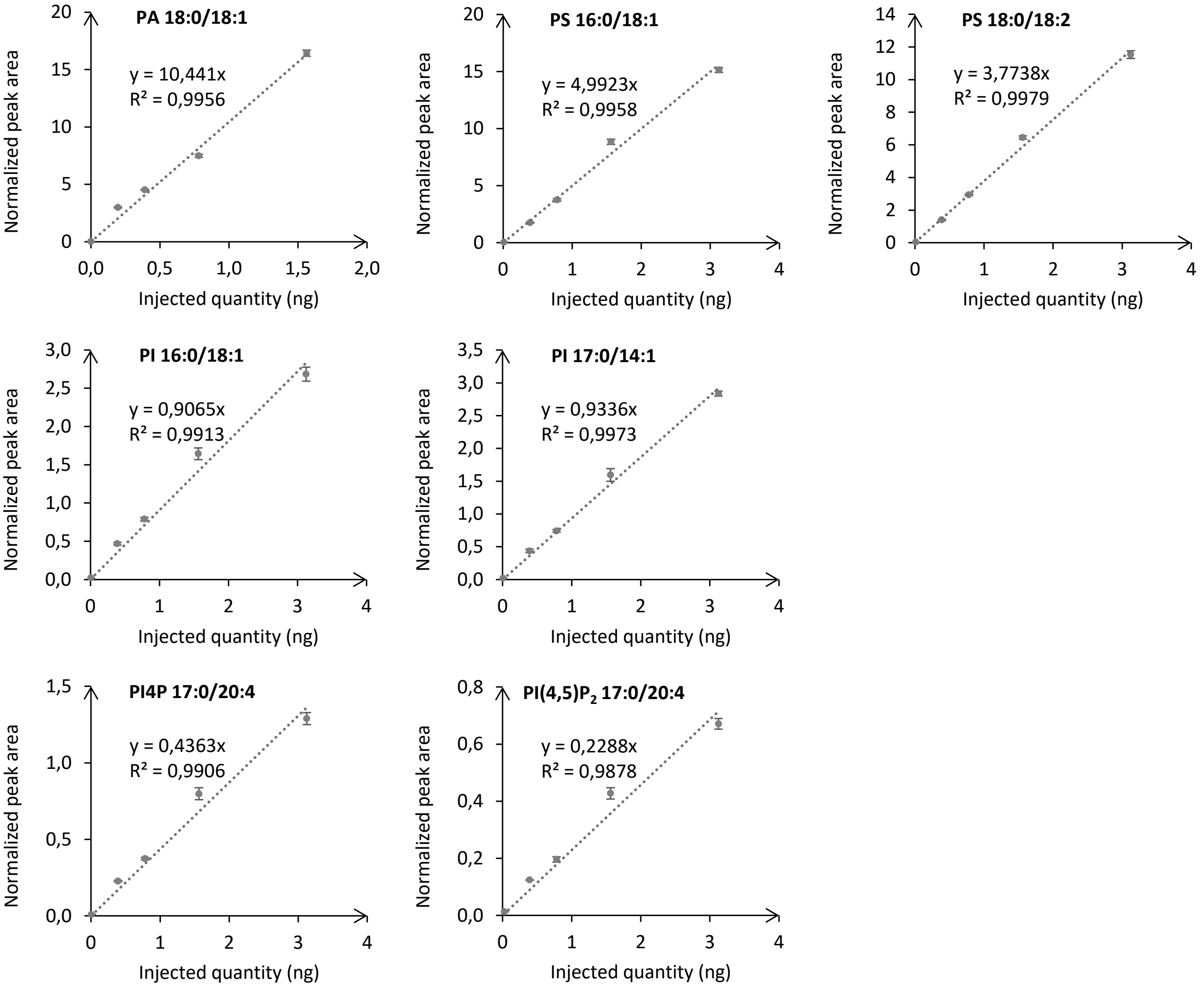


**Figure S1.** Presentation of the calibration curves for methylated anionic phospholipids species. Each point corresponds to the mean of 3 repetitions.

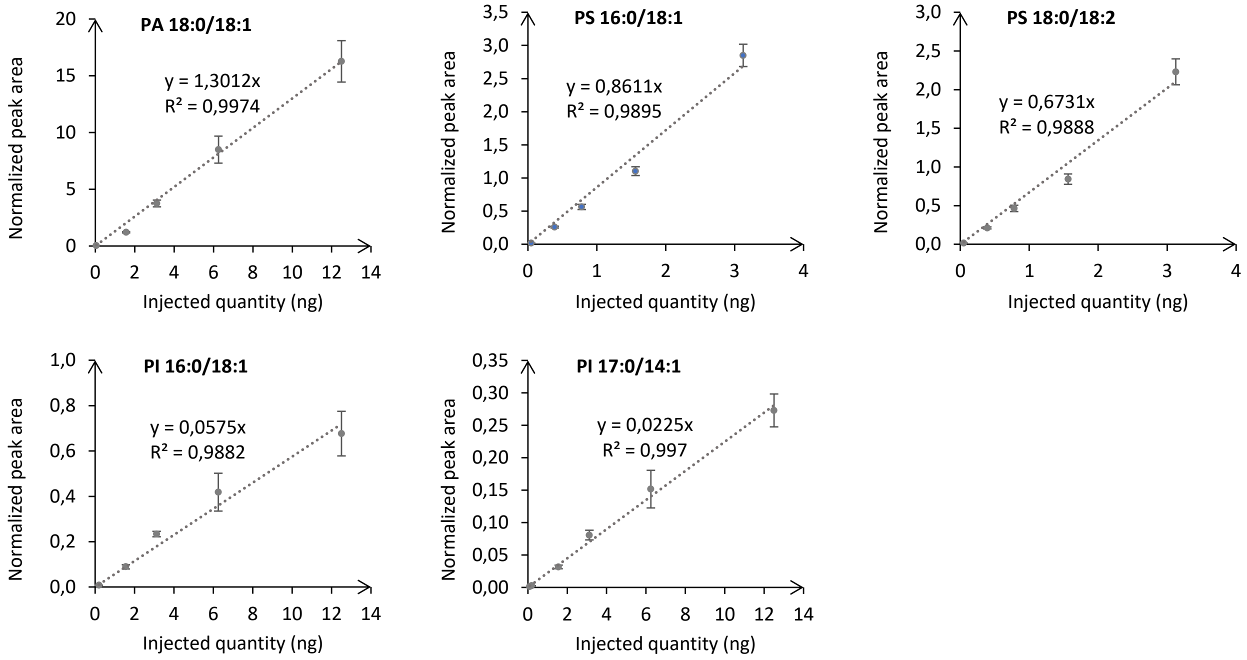


**Figure S2.** Presentation of the calibration curves for non-methylated anionic phospholipids species. Each point corresponds to the mean of 3 repetitions. Of note, no signal was detected for non-methylated PIP and $PIP_2$ lipids.

**a**

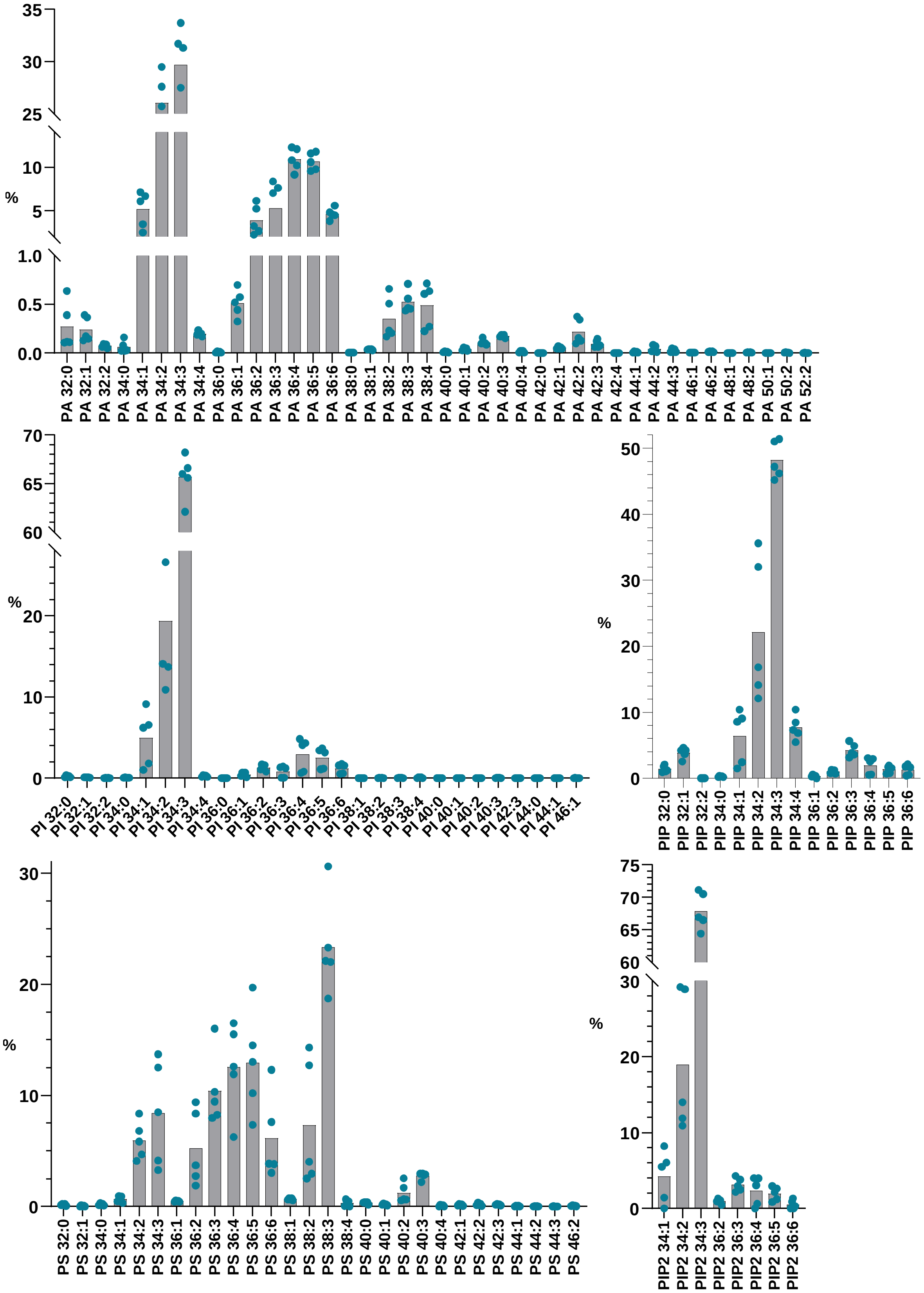

%
35
30
25
10
5
1.0
0.5
0.0
PA 32:0
PA 32:1
PA 32:2
PA 34:0
PA 34:1
PA 34:2
PA 34:3
PA 34:4
PA 36:0
PA 36:1
PA 36:2
PA 36:3
PA 36:4
PA 36:5
PA 36:6
PA 38:0
PA 38:1
PA 38:2
PA 38:3
PA 38:4
PA 40:0
PA 40:1
PA 40:2
PA 40:3
PA 40:4
PA 42:0
PA 42:1
PA 42:2
PA 42:3
PA 42:4
PA 44:1
PA 44:2
PA 44:3
PA 46:1
PA 46:2
PA 48:1
PA 48:2
PA 50:1
PA 50:2
PA 52:2
%
70
65
60
20
10
0
PI 32:0
PI 32:1
PI 32:2
PI 34:0
PI 34:1
PI 34:2
PI 34:3
PI 34:4
PI 36:0
PI 36:1
PI 36:2
PI 36:3
PI 36:4
PI 36:5
PI 36:6
PI 38:1
PI 38:2
PI 38:3
PI 38:4
PI 40:0
PI 40:1
PI 40:2
PI 40:3
PI 42:3
PI 44:0
PI 44:1
PI 46:1
%
50
40
30
20
10
0
PIP 32:0
PIP 32:1
PIP 32:2
PIP 34:0
PIP 34:1
PIP 34:2
PIP 34:3
PIP 34:4
PIP 36:1
PIP 36:2
PIP 36:3
PIP 36:4
PIP 36:5
PIP 36:6
%
30
20
10
0
PS 32:0
PS 32:1
PS 34:0
PS 34:1
PS 34:2
PS 34:3
PS 36:1
PS 36:2
PS 36:3
PS 36:4
PS 36:5
PS 36:6
PS 38:1
PS 38:2
PS 38:3
PS 38:4
PS 40:0
PS 40:1
PS 40:2
PS 40:3
PS 40:4
PS 42:1
PS 42:2
PS 42:3
PS 44:1
PS 44:2
PS 44:3
PS 46:2
%
75
70
65
60
30
20
10
0
PIP2 34:1
PIP2 34:2
PIP2 34:3
PIP2 36:2
PIP2 36:3
PIP2 36:4
PIP2 36:5
PIP2 36:6

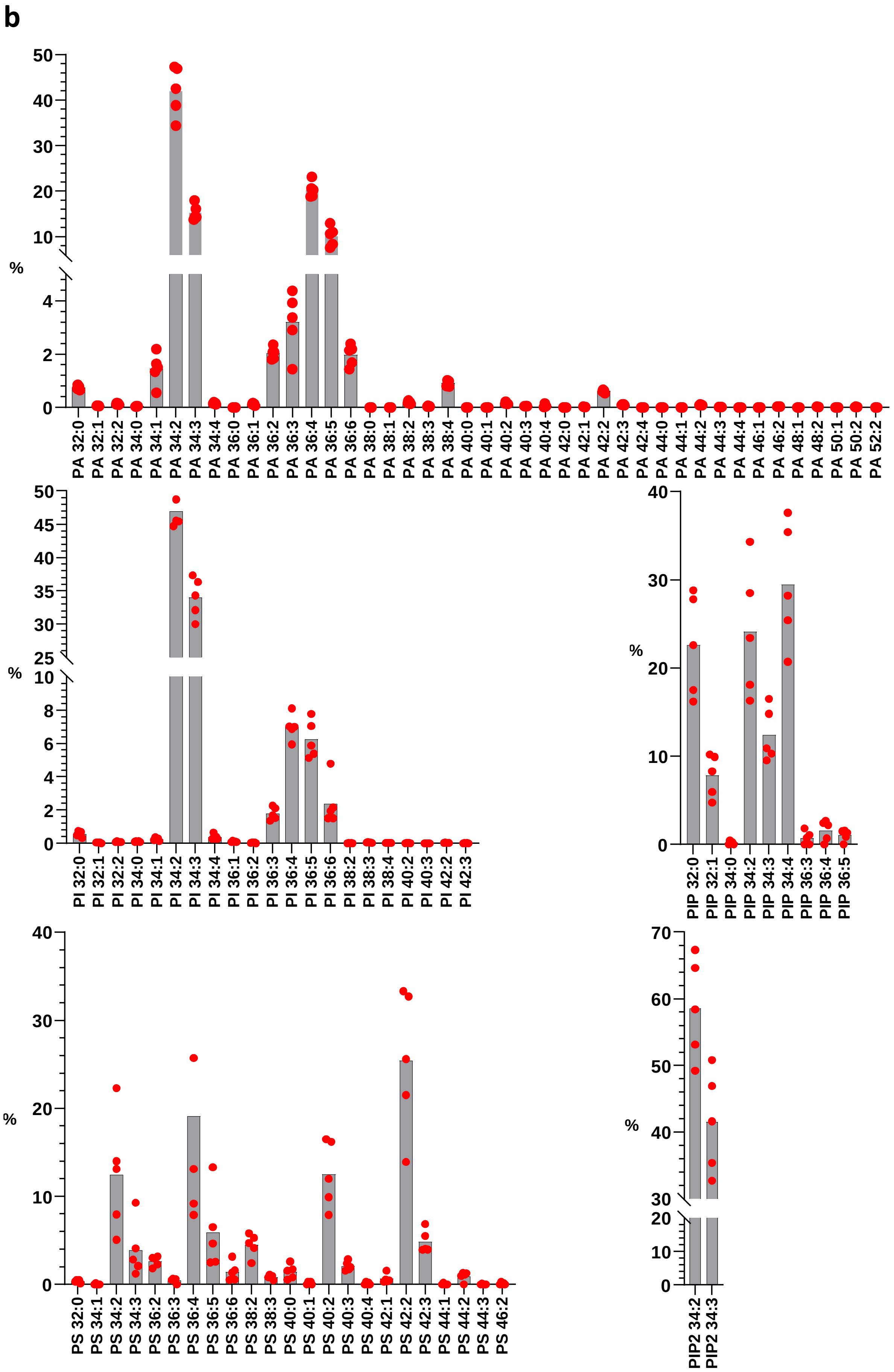


**Figure S3.** Composition of anionic phospholipids in *Nicotiana benthamiana* leaves **(a)** and *Zea mays* leaves **(b)**. Data are expressed in %, n = 5 biological replicates, the dots show the dispersion of data.

**a**

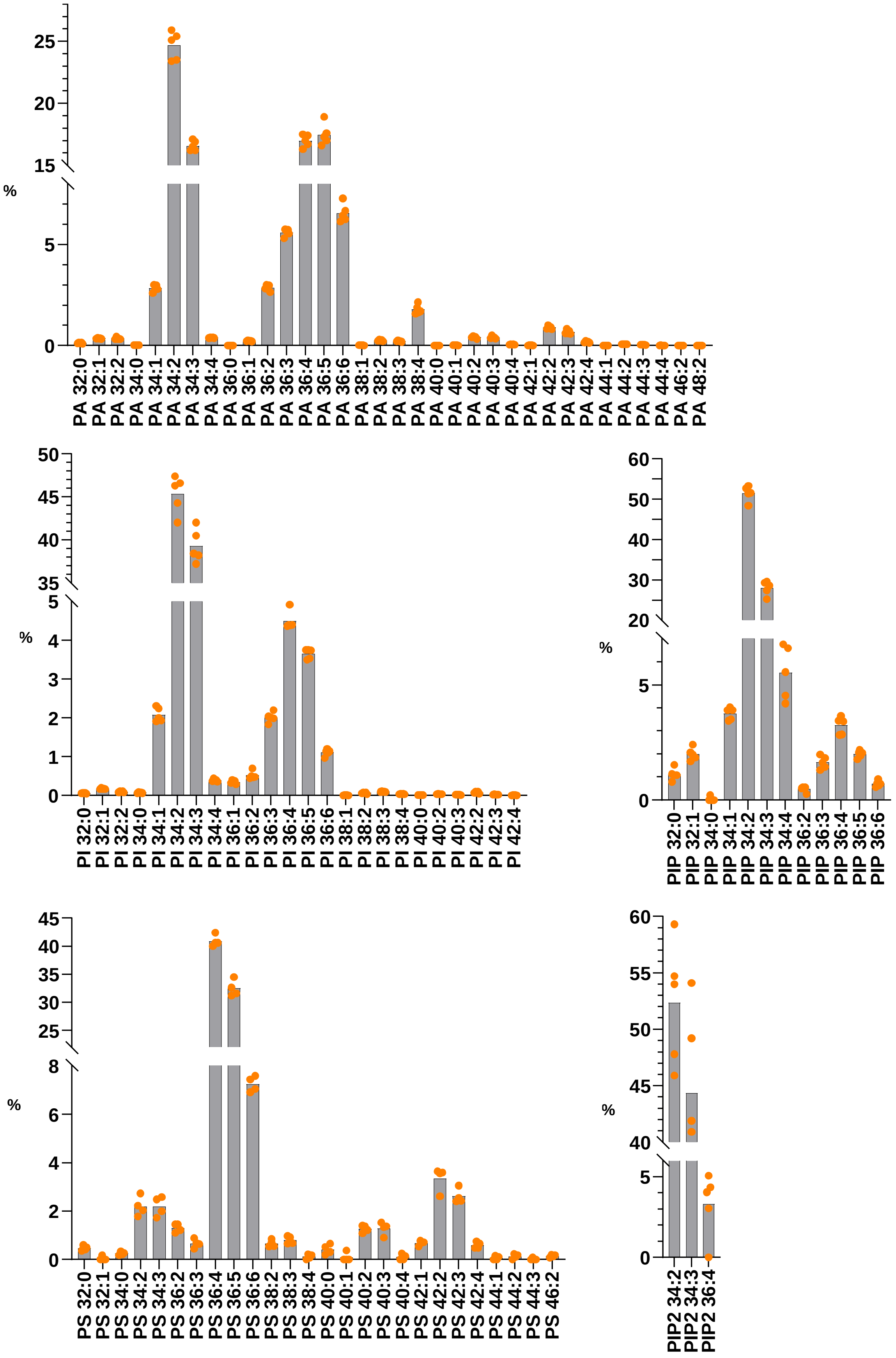

%
25
20
15
5
0
PA 32:0
PA 32:1
PA 32:2
PA 34:0
PA 34:1
PA 34:2
PA 34:3
PA 34:4
PA 36:0
PA 36:1
PA 36:2
PA 36:3
PA 36:4
PA 36:5
PA 36:6
PA 38:1
PA 38:2
PA 38:3
PA 38:4
PA 40:0
PA 40:1
PA 40:2
PA 40:3
PA 40:4
PA 42:1
PA 42:2
PA 42:3
PA 42:4
PA 44:1
PA 44:2
PA 44:3
PA 44:4
PA 46:2
PA 48:2
%
50
45
40
35
5
4
3
2
1
0
PI 32:0
PI 32:1
PI 32:2
PI 34:0
PI 34:1
PI 34:2
PI 34:3
PI 34:4
PI 36:1
PI 36:2
PI 36:3
PI 36:4
PI 36:5
PI 36:6
PI 38:1
PI 38:2
PI 38:3
PI 38:4
PI 40:0
PI 40:2
PI 40:3
PI 42:2
PI 42:3
PI 42:4
%
60
50
40
30
20
5
0
PIP 32:0
PIP 32:1
PIP 34:0
PIP 34:1
PIP 34:2
PIP 34:3
PIP 34:4
PIP 36:2
PIP 36:3
PIP 36:4
PIP 36:5
PIP 36:6
%
45
40
35
30
25
8
6
4
2
0
PS 32:0
PS 32:1
PS 34:0
PS 34:2
PS 34:3
PS 36:2
PS 36:3
PS 36:4
PS 36:5
PS 36:6
PS 38:2
PS 38:3
PS 38:4
PS 40:0
PS 40:1
PS 40:2
PS 40:3
PS 40:4
PS 42:1
PS 42:2
PS 42:3
PS 42:4
PS 44:1
PS 44:2
PS 44:3
PS 46:2
%
60
55
50
45
40
5
0
PIP2 34:2
PIP2 34:3
PIP2 36:4

**b**

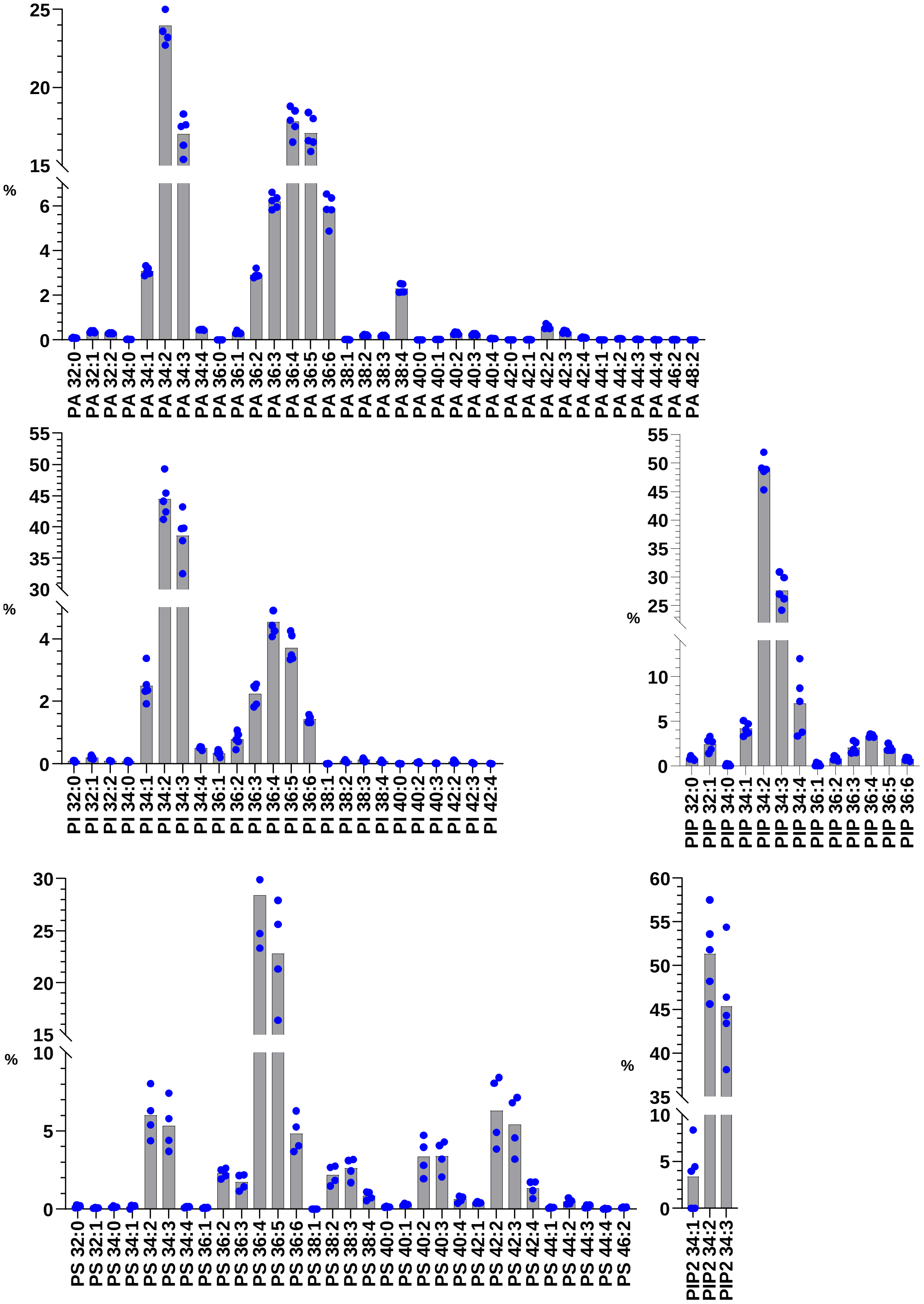

25
20
15
%
6
4
2
0
PA 32:0
PA 32:1
PA 32:2
PA 34:0
PA 34:1
PA 34:2
PA 34:3
PA 34:4
PA 36:0
PA 36:1
PA 36:2
PA 36:3
PA 36:4
PA 36:5
PA 36:6
PA 38:1
PA 38:2
PA 38:3
PA 38:4
PA 40:0
PA 40:1
PA 40:2
PA 40:3
PA 40:4
PA 42:0
PA 42:1
PA 42:2
PA 42:3
PA 42:4
PA 44:1
PA 44:2
PA 44:3
PA 44:4
PA 46:2
PA 48:2
55
50
45
40
35
30
%
4
2
0
PI 32:0
PI 32:1
PI 32:2
PI 34:0
PI 34:1
PI 34:2
PI 34:3
PI 34:4
PI 36:1
PI 36:2
PI 36:3
PI 36:4
PI 36:5
PI 36:6
PI 38:1
PI 38:2
PI 38:3
PI 38:4
PI 40:0
PI 40:2
PI 40:3
PI 42:2
PI 42:3
PI 42:4
55
50
45
40
35
30
25
%
10
5
0
PIP 32:0
PIP 32:1
PIP 34:0
PIP 34:1
PIP 34:2
PIP 34:3
PIP 34:4
PIP 36:1
PIP 36:2
PIP 36:3
PIP 36:4
PIP 36:5
PIP 36:6
30
25
20
15
%
10
5
0
PS 32:0
PS 32:1
PS 34:0
PS 34:1
PS 34:2
PS 34:3
PS 34:4
PS 36:1
PS 36:2
PS 36:3
PS 36:4
PS 36:5
PS 36:6
PS 38:1
PS 38:2
PS 38:3
PS 38:4
PS 40:0
PS 40:1
PS 40:2
PS 40:3
PS 40:4
PS 42:1
PS 42:2
PS 42:3
PS 42:4
PS 44:1
PS 44:2
PS 44:3
PS 44:4
PS 46:2
60
55
50
45
40
35
%
10
5
0
PIP2 34:1
PIP2 34:2
PIP2 34:3

c

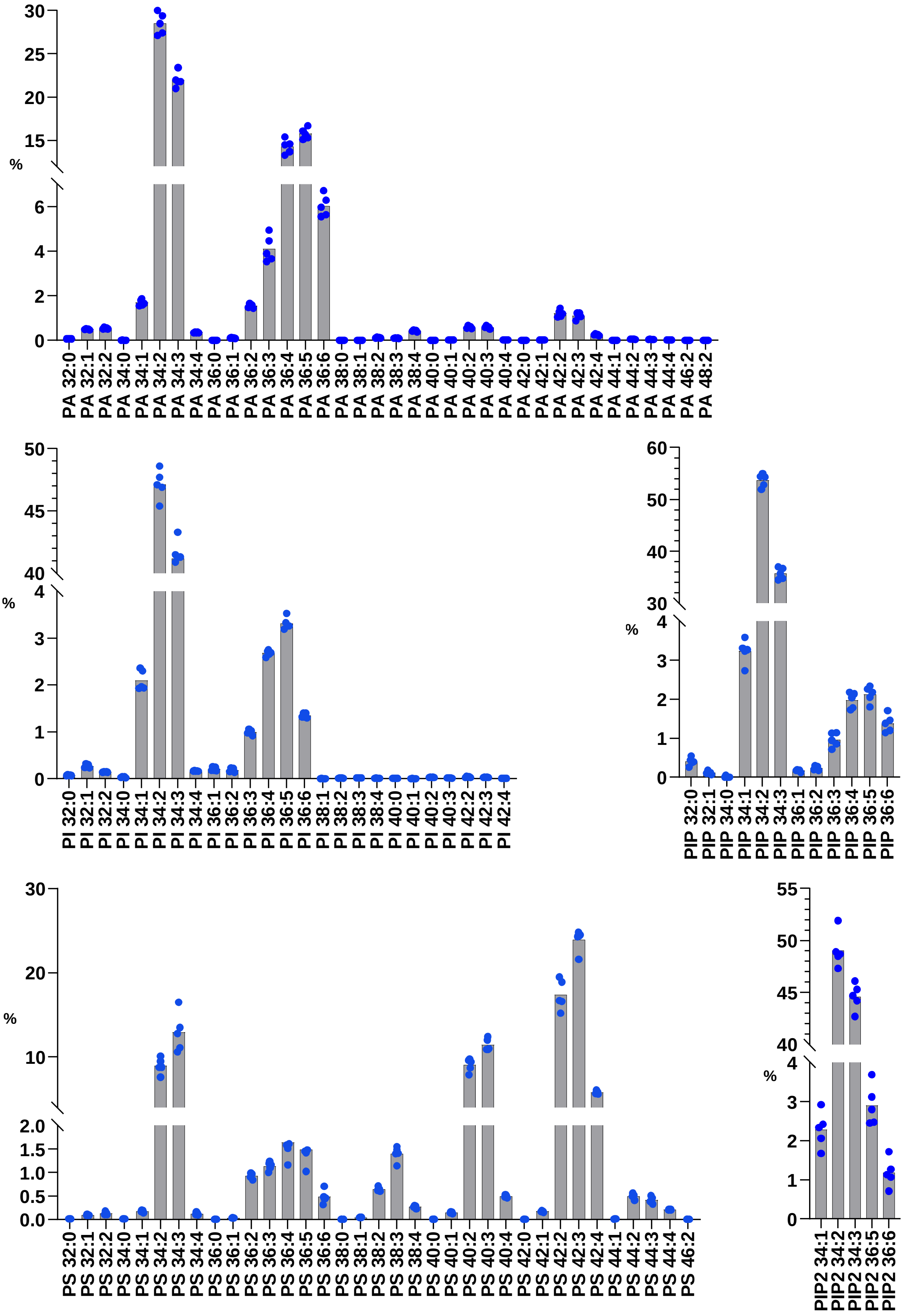
%
30
25
20
15
6
4
2
0
PA 32:0
PA 32:1
PA 32:2
PA 34:0
PA 34:1
PA 34:2
PA 34:3
PA 34:4
PA 36:0
PA 36:1
PA 36:2
PA 36:3
PA 36:4
PA 36:5
PA 36:6
PA 38:0
PA 38:1
PA 38:2
PA 38:3
PA 38:4
PA 40:0
PA 40:1
PA 40:2
PA 40:3
PA 40:4
PA 42:0
PA 42:1
PA 42:2
PA 42:3
PA 42:4
PA 44:1
PA 44:2
PA 44:3
PA 44:4
PA 46:2
PA 48:2
%
50
45
40
4
3
2
1
0
PI 32:0
PI 32:1
PI 32:2
PI 34:0
PI 34:1
PI 34:2
PI 34:3
PI 34:4
PI 36:1
PI 36:2
PI 36:3
PI 36:4
PI 36:5
PI 36:6
PI 38:1
PI 38:2
PI 38:3
PI 38:4
PI 40:0
PI 40:1
PI 40:2
PI 40:3
PI 42:2
PI 42:3
PI 42:4
%
60
50
40
30
4
3
2
1
0
PIP 32:0
PIP 32:1
PIP 34:0
PIP 34:1
PIP 34:2
PIP 34:3
PIP 36:1
PIP 36:2
PIP 36:3
PIP 36:4
PIP 36:5
PIP 36:6
%
30
20
10
2.0
1.5
1.0
0.5
0.0
PS 32:0
PS 32:1
PS 32:2
PS 34:0
PS 34:1
PS 34:2
PS 34:3
PS 34:4
PS 36:0
PS 36:1
PS 36:2
PS 36:3
PS 36:4
PS 36:5
PS 36:6
PS 38:0
PS 38:1
PS 38:2
PS 38:3
PS 38:4
PS 40:0
PS 40:1
PS 40:2
PS 40:3
PS 40:4
PS 42:0
PS 42:1
PS 42:2
PS 42:3
PS 42:4
PS 44:1
PS 44:2
PS 44:3
PS 44:4
PS 46:2
%
55
50
45
40
4
3
2
1
0
PIP2 34:1
PIP2 34:2
PIP2 34:3
PIP2 36:5
PIP2 36:6

**d**

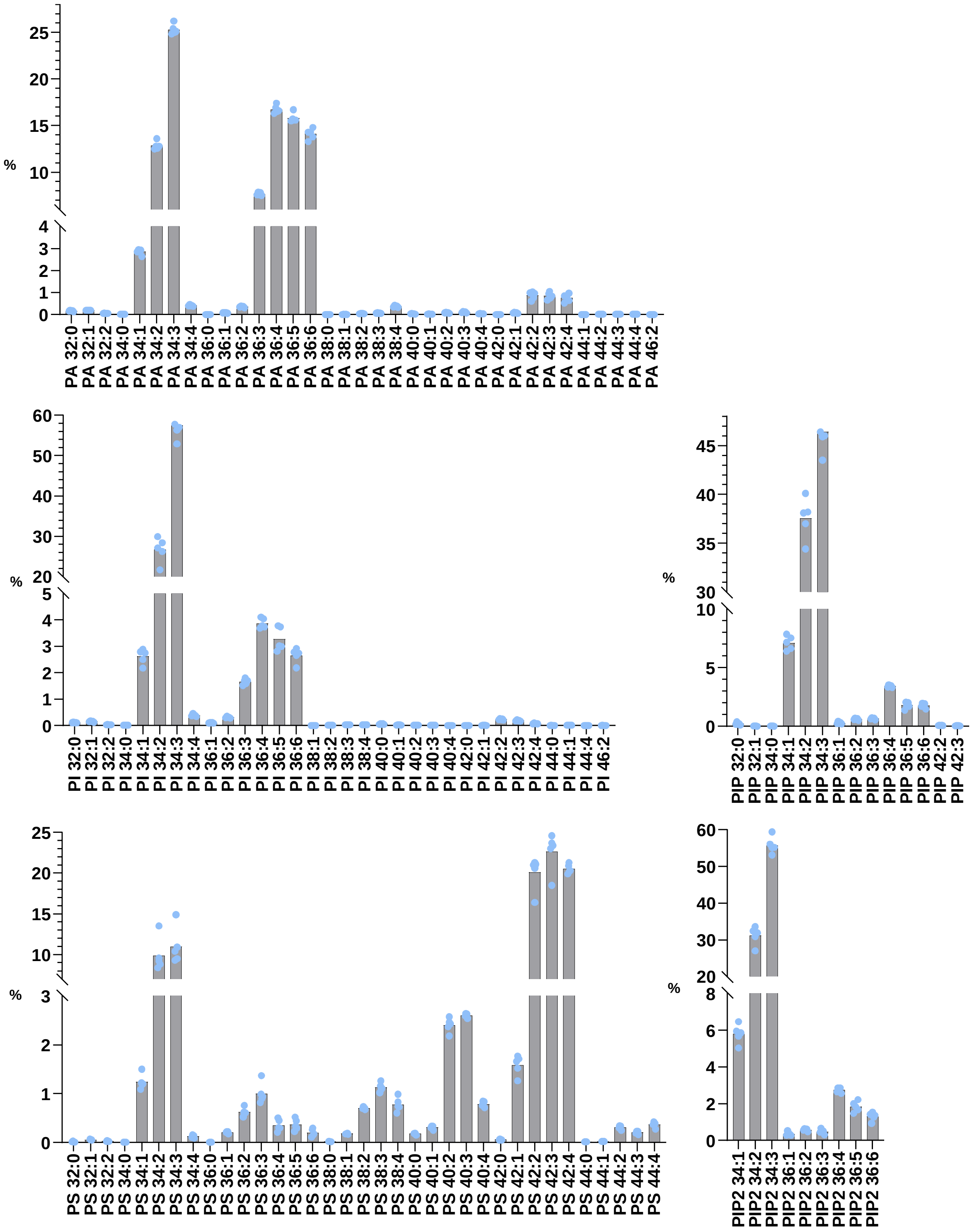


**Figure S4.** Composition of *Arabidopsis thaliana* anionic phospholipids in 7-days old seedlings **(a)**, aerial part (cotyledons and primary leaves) **(b)**, roots **(c)** and suspension cells 10-days after sub-culturing **(d)**. Data are expressed in %, n = 5 biological replicates, the dots show the dispersion of data.

**Table S1.** For a reaction time of 10 min at 23°C, presentation of the percentage of the investigated anionic lipid species methylated various times (n=3).

| **Investigated lipid species** | **Percentage (%) of anionic phospholipids with increasing additional methyl groups following methylation** | | | | | | | |
|---|---|---|---|---|---|---|---|---|
| | **0** | **1** | **2** | **3** | **4** | **5** | **6** | **7** |
| **PA 32:0** | 0,0±0,0 | 0,0±0,0 | 98,5±0,8 | 0,7±0,3 | 0,8±0,7 | | | |
| **PA 32:1** | 0,0±0,0 | 1,0±0,4 | 95,1±1,1 | 2,1±0,8 | 1,8±0,6 | | | |
| **PA 32:2** | 0,0±0,0 | 7,0±2,5 | 92,1±2,4 | 0,9±0,1 | 0,0±0,0 | | | |
| **PA 34:0** | 0,0±0,0 | 0,0±0,0 | 98,8±1,2 | 0,0±0,0 | 1,2±1,2 | | | |
| **PA 34:1** | 0,0±0,0 | 0,7±0,3 | 98,0±0,1 | 0,8±0,2 | 0,5±0,2 | | | |
| **PA 34:2** | 0,0±0,0 | 11,5±4,2 | 87,4±4,5 | 0,6±0,1 | 0,5±0,2 | | | |
| **PA 36:1** | 0,0±0,0 | 0,4±0,2 | 97,5±0,6 | 1,4±0,5 | 0,7±0,3 | | | |
| **PA 36:2** | 0,0±0,0 | 3,4±0,4 | 94,9±0,4 | 1,1±0,4 | 0,6±0,3 | | | |
| **PA 38:4** | 0,0±0,0 | 0,3±0,0 | 98,0±0,6 | 1,0±0,4 | 0,7±0,3 | | | |
| **PA 38:5** | 0,0±0,0 | 0,0±0,0 | 97,9±0,6 | 1,5±0,2 | 0,6±0,7 | | | |
| **PA 38:6** | 0,0±0,0 | 0,0±0,0 | 99,5±0,1 | 0,2±0,0 | 0,3±0,1 | | | |
| **PA 40:6** | 0,0±0,0 | 0,0±0,0 | 97,5±0,4 | 1,5±0,3 | 1,0±0,1 | | | |
| **Mean** | **0,0±0,0** | **2,0±3,6** | **96,3±3,5** | **1,0±0,6** | **0,7±0,5** | | | |
| **PI 34:1** | 0,0±0,0 | 78,5±2,6 | 18,2±2,7 | 3,4±0,4 | 0,0±0,0 | | | |
| **PI 34:2** | 0,0±0,0 | 86,1±3,5 | 12,9±3,3 | 1,0±0,2 | 0,0±0,0 | | | |
| **PI 34:3** | 0,0±0,0 | 89,7±2,1 | 9,1±2,2 | 1,2±0,1 | 0,0±0,0 | | | |
| **PI 36:1** | 0,0±0,0 | 78,4±7,9 | 20,4±6,9 | 1,2±1,0 | 0,0±0,0 | | | |
| **PI 36:2** | 0,4±0,2 | 78,2±5,6 | 18,2±4,8 | 3,2±1,2 | 0,0±0,0 | | | |
| **PI 36:3** | 0,1±0,1 | 81,1±6,5 | 17,0±6,0 | 1,9±0,5 | 0,0±0,0 | | | |
| **PI 36:4** | 0,1±0,1 | 85,6±3,5 | 13,2±3,8 | 1,2±0,6 | 0,0±0,0 | | | |
| **PI 36:5** | 0,0±0,0 | 77,9±2,1 | 20,2±1,7 | 2,0±0,5 | 0,0±0,0 | | | |
| **Mean** | **0,1±0,1** | **81,9±4,6** | **16,1±4,0** | **1,9±0,9** | **0,0±0,0** | | | |
| **PS 34:1** | 0,0±0,0 | 0,1±0,0 | 81,4±1,0 | 18,5±1,0 | 0,0±0,0 | | | |
| **PS 34:2** | 0,0±0,0 | 0,1±0,0 | 79,2±1,5 | 20,7±1,5 | 0,1±0,0 | | | |
| **PS 34:3** | 0,0±0,0 | 0,2±0,0 | 81,8±1,7 | 18,0±1,7 | 0,1±0,0 | | | |
| **PS 36:1** | 0,0±0,0 | 3,2±0,7 | 87,9±1,4 | 8,4±1,1 | 0,4±0,1 | | | |
| **PS 36:2** | 0,0±0,0 | 0,4±0,1 | 88,6±1,2 | 10,9±1,2 | 0,1±0,0 | | | |
| **PS 36:3** | 0,0±0,0 | 0,1±0,0 | 84,0±0,5 | 15,8±0,4 | 0,1±0,1 | | | |
| **PS 36:4** | 0,0±0,0 | 0,1±0,0 | 75,7±2,0 | 24,2±2,0 | 0,0±0,0 | | | |
| **PS 36:5** | 0,0±0,0 | 0,1±0,0 | 79,8±1,6 | 19,9±1,6 | 0,2±0,0 | | | |
| **PS 36:6** | 0,0±0,0 | 0,0±0,0 | 74,5±6,7 | 25,1±6,6 | 0,3±0,1 | | | |
| **Mean** | **0,0±0,0** | **0,5±1,0** | **81,4±4,9** | **18,0±5,5** | **0,1±0,1** | | | |
| **PIP 32:0** | 0,0±0,0 | 0,0±0,0 | 0,0±0,0 | 56,9±3,2 | 33,7±3,0 | 9,3±2,9 | 0,0±0,0 | |
| **PIP 34:0** | 0,0±0,0 | 0,4±0,7 | 0,0±0,0 | 57,8±5,3 | 33,1±3,9 | 8,7±1,8 | 0,0±0,0 | |
| **PIP 34:1** | 0,0±0,0 | 0,0±0,0 | 0,0±0,0 | 59,1±1,4 | 30,5±2,2 | 9,7±1,2 | 0,7±0,2 | |
| **PIP 36:1** | 0,0±0,0 | 0,0±0,0 | 0,1±0,1 | 63,6±0,3 | 30,0±0,9 | 5,7±1,0 | 0,6±0,0 | |
| **PIP 36:2** | 0,0±0,0 | 0,0±0,0 | 0,0±0,0 | 57,5±2,6 | 34,3±1,3 | 7,6±1,2 | 0,6±0,1 | |
| **PIP 36:3** | 0,0±0,0 | 0,0±0,0 | 0,0±0,0 | 58,5±6,3 | 33,8±4,1 | 7,7±2,8 | 0,0±0,0 | |
| **PIP 36:4** | 0,0±0,0 | 0,0±0,0 | 0,0±0,0 | 73,8±2,2 | 21,1±1,3 | 4,6±0,9 | 0,5±0,1 | |
| **PIP 38:2** | 0,0±0,0 | 0,0±0,0 | 0,0±0,0 | 62,2±4,4 | 30,5±2,7 | 6,6±0,7 | 0,7±1,1 | |
| **PIP 38:3** | 0,0±0,0 | 0,0±0,0 | 0,0±0,0 | 63,2±5,7 | 30,8±4,9 | 6,0±2,1 | 0,0±0,0 | |
| **PIP 38:4** | 0,0±0,0 | 0,0±0,0 | 0,3±0,0 | 52,0±4,1 | 36,5±2,5 | 10,4±0,9 | 0,8±0,7 | |
| **PIP 38:5** | 0,0±0,0 | 0,0±0,0 | 0,0±0,0 | 71,7±1,5 | 21,5±0,7 | 6,2±0,8 | 0,6±0,1 | |
| **PIP 40:4** | 0,0±0,0 | 0,0±0,0 | 0,0±0,0 | 63,9±2,3 | 29,3±1,8 | 6,3±0,5 | 0,5±0,1 | |
| **Mean** | **0,0±0,0** | **0,0±0,1** | **0,0±0,1** | **61,7±6,2** | **30,4±4,8** | **7,4±1,8** | **0,4±0,3** | |
| **PI(4,5)P$_2$ 32:0** | 0,0±0,0 | 0,0±0,0 | 0,0±0,0 | 0,0±0,0 | 0,0±0,0 | 64,4±4,9 | 30,7±4,0 | 4,9±0,9 |
| **PI(4,5)P$_2$ 34:0** | 0,0±0,0 | 0,0±0,0 | 0,0±0,0 | 0,0±0,0 | 0,0±0,0 | 73,3±2,8 | 23,2±2,0 | 3,5±1,2 |
| **PI(4,5)P$_2$ 34:1** | 0,0±0,0 | 0,0±0,0 | 0,0±0,0 | 0,0±0,0 | 0,2±0,2 | 69,6±2,0 | 26,7±1,6 | 3,5±0,4 |
| **PI(4,5)P$_2$ 36:1** | 0,0±0,0 | 0,0±0,0 | 0,0±0,0 | 0,0±0,0 | 0,3±0,1 | 71,0±1,4 | 25,7±1,0 | 2,9±0,4 |
| **PI(4,5)P$_2$ 36:2** | 0,0±0,0 | 0,0±0,0 | 0,0±0,0 | 0,0±0,0 | 0,2±0,0 | 67,3±1,2 | 25,4±0,9 | 3,2±0,4 |
| **PI(4,5)P$_2$ 36:3** | 0,0±0,0 | 0,0±0,0 | 0,0±0,0 | 0,0±0,0 | 0,3±0,4 | 66,8±4,0 | 28,9±1,7 | 4,1±2,2 |
| **PI(4,5)P$_2$ 36:4** | 0,0±0,0 | 0,0±0,0 | 0,0±0,0 | 0,0±0,0 | 0,0±0,0 | 72,6±0,9 | 25,0±0,7 | 2,4±0,5 |
| **PI(4,5)P$_2$ 38:3** | 0,0±0,1 | 0,0±0,0 | 0,0±0,0 | 0,0±0,0 | 0,0±0,0 | 72,1±1,4 | 24,3±1,6 | 3,5±0,2 |
| **PI(4,5)P$_2$ 38:4** | 0,0±0,0 | 0,0±0,0 | 0,0±0,0 | 0,0±0,0 | 0,4±0,0 | 71,7±2,5 | 24,6±2,6 | 3,3±0,3 |
| **PI(4,5)P$_2$ 38:5** | 0,0±0,0 | 0,0±0,0 | 0,0±0,0 | 0,0±0,0 | 0,4±0,1 | 72,8±0,6 | 24,4±0,8 | 2,4±0,3 |
| **PI(4,5)P$_2$ 40:4** | 0,0±0,0 | 0,0±0,0 | 0,0±0,0 | 0,0±0,0 | 0,4±0,1 | 79,3±0,7 | 17,6±0,8 | 2,7±0,6 |

| | | | | | | | | |
|---|---|---|---|---|---|---|---|---|
| **PI(4,5)$P_2$ 40:5** | 0,0±0,0 | 0,0±0,0 | 0,0±0,0 | 0,0±0,0 | 0,0±0,0 | 67,6±1,1 | 29,5±1,5 | 2,9±0,5 |
| **Mean** | **0,0±0,0** | **0,0±0,0** | **0,0±0,0** | **0,0±0,0** | **0,2±0,2** | **70,7±3,9** | **25,5±3,4** | **3,3±0,7** |

**Table S2.** List of MRM information (Q1 and Q3 mass, DP, CE and CXP) used for the analysis of anionic PL commercial mixes in the context of the optimization of chromatographic separation and of ionization temperature.

| Analyte | Q1 Mass (Da) | Q3 mass (Da) | DP (V) | CE (V) | CXP (V) |
|---|---|---|---|---|---|
| PI 34:2 | 866,6 | 575,5 | 10 | 31 | 25 |
| PI 36:2 | 894,6 | 603,5 | 10 | 31 | 25 |
| PI 34:1 | 868,6 | 577,5 | 10 | 31 | 25 |
| PI 34:3 | 864,5 | 573,5 | 10 | 31 | 25 |
| PI 36:1 | 896,6 | 605,6 | 10 | 31 | 25 |
| PI 36:3 | 892,6 | 601,5 | 10 | 31 | 25 |
| PI 36:4 | 890,6 | 599,5 | 10 | 31 | 25 |
| PI 36:5 | 888,5 | 597,5 | 10 | 31 | 25 |
| PA 34:1 | 720,6 | 577,5 | 10 | 26 | 19 |
| PA 36:1 | 748,6 | 605,6 | 10 | 26 | 19 |
| PA 36:2 | 746,6 | 603,5 | 10 | 26 | 19 |
| PA 38:4 | 770,5 | 627,5 | 10 | 26 | 19 |
| PA 38:5 | 768,5 | 625,5 | 10 | 26 | 19 |
| PA 38:6 | 766,6 | 623,5 | 10 | 26 | 19 |
| PA 40:6 | 794,6 | 651,5 | 10 | 26 | 19 |
| PA 32:0 | 694,5 | 551,5 | 10 | 26 | 19 |
| PA 32:1 | 692,6 | 549,5 | 10 | 26 | 19 |
| PA 32:2 | 690,6 | 547,5 | 10 | 26 | 19 |
| PA 34:0 | 722,6 | 579,5 | 10 | 26 | 19 |
| PA 34:2 | 718,5 | 575,5 | 10 | 26 | 19 |
| PS 34:2 | 788,5 | 575,5 | 10 | 34 | 24 |
| PS 36:4 | 812,5 | 599,5 | 10 | 34 | 24 |
| PS 34:1 | 790,5 | 577,5 | 10 | 34 | 24 |
| PS 34:3 | 786,5 | 573,5 | 10 | 34 | 24 |
| PS 36:1 | 818,6 | 605,6 | 10 | 34 | 24 |
| PS 36:2 | 816,5 | 603,5 | 10 | 34 | 24 |
| PS 36:3 | 814,5 | 601,5 | 10 | 34 | 24 |
| PS 36:5 | 810,5 | 597,5 | 10 | 34 | 24 |
| PS 36:6 | 808,5 | 595,5 | 10 | 34 | 24 |
| PI4P 32:0 | 933,6 | 551,5 | 10 | 32 | 24 |
| PI4P 34:0 | 961,6 | 579,5 | 10 | 32 | 24 |
| PI4P 34:1 | 959,6 | 577,5 | 10 | 32 | 24 |
| PI4P 36:1 | 987,6 | 605,6 | 10 | 32 | 24 |
| PI4P 36:2 | 985,6 | 603,5 | 10 | 32 | 24 |
| PI4P 36:3 | 983,6 | 601,5 | 10 | 32 | 24 |
| PI4P 36:4 | 981,6 | 599,5 | 10 | 32 | 24 |
| PI4P 38:2 | 1013,6 | 631,6 | 10 | 32 | 24 |
| PI4P 38:3 | 1011,6 | 629,6 | 10 | 32 | 24 |
| PI4P 38:4 | 1009,6 | 627,5 | 10 | 32 | 24 |
| PI4P 38:5 | 1007,6 | 625,5 | 10 | 32 | 24 |
| PI4P 40:4 | 1037,6 | 655,6 | 10 | 32 | 24 |
| PI(4,5)$P_2$ 32:0 | 1041,6 | 551,5 | 10 | 45 | 32 |
| PI(4,5)$P_2$ 34:0 | 1069,6 | 579,5 | 10 | 45 | 32 |
| PI(4,5)$P_2$ 34:1 | 1067,6 | 577,5 | 10 | 45 | 32 |
| PI(4,5)$P_2$ 36:1 | 1095,6 | 605,6 | 10 | 45 | 32 |
| PI(4,5)$P_2$ 36:2 | 1093,6 | 603,5 | 10 | 45 | 32 |
| PI(4,5)$P_2$ 36:3 | 1091,6 | 601,5 | 10 | 45 | 32 |
| PI(4,5)$P_2$ 36:4 | 1089,6 | 599,5 | 10 | 45 | 32 |
| PI(4,5)$P_2$ 38:3 | 1119,6 | 629,6 | 10 | 45 | 32 |
| PI(4,5)$P_2$ 38:4 | 1117,6 | 627,5 | 10 | 45 | 32 |
| PI(4,5)$P_2$ 38:5 | 1115,6 | 625,5 | 10 | 45 | 32 |
| PI(4,5)$P_2$ 40:4 | 1145,6 | 655,6 | 10 | 45 | 32 |
| PI(4,5)$P_2$ 40:5 | 1143,6 | 653,6 | 10 | 45 | 32 |

**Table S3.** List of MRM information (Q1 and Q3 mass, DP, CE and CXP) used for the analysis of anionic PL commercial mixes in the context of the optimization of the methylation reaction conditions.

| Analyte | Q1 Mass (Da) | Q3 mass (Da) | DP (V) | CE (V) | CXP (V) |
|---|---|---|---|---|---|
| PI 34:2 Me0 | 852,6 | 575,5 | 10 | 31 | 25 |
| PI 34:2 Me1 | 866,6 | 575,5 | 10 | 31 | 25 |
| PI 34:2 Me2 | 880,6 | 575,5 | 10 | 31 | 25 |
| PI 34:2 Me3 | 894,6 | 575,5 | 10 | 31 | 25 |
| PI 34:2 Me4 | 908,6 | 575,5 | 10 | 31 | 25 |
| PI 36:2 Me0 | 880,6 | 603,5 | 10 | 31 | 25 |
| PI 36:2 Me1 | 894,6 | 603,5 | 10 | 31 | 25 |
| PI 36:2 Me2 | 908,6 | 603,5 | 10 | 31 | 25 |
| PI 36:2 Me3 | 922,6 | 603,5 | 10 | 31 | 25 |
| PI 36:2 Me4 | 936,7 | 603,5 | 10 | 31 | 25 |
| PI 34:1 Me0 | 854,6 | 577,5 | 10 | 31 | 25 |
| PI 34:1 Me1 | 868,6 | 577,5 | 10 | 31 | 25 |
| PI 34:1 Me2 | 882,6 | 577,5 | 10 | 31 | 25 |
| PI 34:1 Me3 | 896,6 | 577,5 | 10 | 31 | 25 |
| PI 34:1 Me4 | 910,7 | 577,5 | 10 | 31 | 25 |
| PI 34:3 Me0 | 850,5 | 573,5 | 10 | 31 | 25 |
| PI 34:3 Me1 | 864,5 | 573,5 | 10 | 31 | 25 |
| PI 34:3 Me2 | 878,5 | 573,5 | 10 | 31 | 25 |
| PI 34:3 Me3 | 892,5 | 573,5 | 10 | 31 | 25 |
| PI 34:3 Me4 | 906,6 | 573,5 | 10 | 31 | 25 |
| PI 36:1 Me0 | 882,6 | 605,6 | 10 | 31 | 25 |
| PI 36:1 Me1 | 896,6 | 605,6 | 10 | 31 | 25 |
| PI 36:1 Me2 | 910,6 | 605,6 | 10 | 31 | 25 |
| PI 36:1 Me3 | 924,6 | 605,6 | 10 | 31 | 25 |
| PI 36:1 Me4 | 938,7 | 605,6 | 10 | 31 | 25 |
| PI 36:3 Me0 | 878,6 | 601,5 | 10 | 31 | 25 |
| PI 36:3 Me1 | 892,6 | 601,5 | 10 | 31 | 25 |
| PI 36:3 Me2 | 906,6 | 601,5 | 10 | 31 | 25 |
| PI 36:3 Me3 | 920,6 | 601,5 | 10 | 31 | 25 |
| PI 36:3 Me4 | 934,7 | 601,5 | 10 | 31 | 25 |
| PI 36:4 Me0 | 876,6 | 599,5 | 10 | 31 | 25 |
| PI 36:4 Me1 | 890,6 | 599,5 | 10 | 31 | 25 |
| PI 36:4 Me2 | 904,6 | 599,5 | 10 | 31 | 25 |
| PI 36:4 Me3 | 918,6 | 599,5 | 10 | 31 | 25 |
| PI 36:4 Me4 | 932,7 | 599,5 | 10 | 31 | 25 |
| PI 36:5 Me0 | 874,5 | 597,5 | 10 | 31 | 25 |
| PI 36:5 Me1 | 888,5 | 597,5 | 10 | 31 | 25 |
| PI 36:5 Me2 | 902,5 | 597,5 | 10 | 31 | 25 |
| PI 36:5 Me3 | 916,5 | 597,5 | 10 | 31 | 25 |
| PI 36:5 Me4 | 930,6 | 597,5 | 10 | 31 | 25 |
| PA 34:1 Me0 | 692,5 | 577,5 | 10 | 26 | 19 |
| PA 34:1 Me1 | 706,5 | 577,5 | 10 | 26 | 19 |
| PA 34:1 Me2 | 720,6 | 577,5 | 10 | 26 | 19 |
| PA 34:1 Me3 | 734,6 | 577,5 | 10 | 26 | 19 |
| PA 34:1 Me4 | 748,6 | 577,5 | 10 | 26 | 19 |
| PA 36:1 Me0 | 720,6 | 605,6 | 10 | 26 | 19 |
| PA 36:1 Me1 | 734,6 | 605,6 | 10 | 26 | 19 |
| PA 36:1 Me2 | 748,6 | 605,6 | 10 | 26 | 19 |
| PA 36:1 Me3 | 762,6 | 605,6 | 10 | 26 | 19 |
| PA 36:1 Me4 | 776,6 | 605,6 | 10 | 26 | 19 |
| PA 36:2 Me0 | 718,5 | 603,5 | 10 | 26 | 19 |
| PA 36:2 Me1 | 732,6 | 603,5 | 10 | 26 | 19 |
| PA 36:2 Me2 | 746,6 | 603,5 | 10 | 26 | 19 |
| PA 36:2 Me3 | 760,6 | 603,5 | 10 | 26 | 19 |
| PA 36:2 Me4 | 774,6 | 603,5 | 10 | 26 | 19 |
| PA 38:4 Me0 | 742,5 | 627,5 | 10 | 26 | 19 |
| PA 38:4 Me1 | 756,5 | 627,5 | 10 | 26 | 19 |
| PA 38:4 Me2 | 770,5 | 627,5 | 10 | 26 | 19 |

| | | | | | |
|---|---|---|---|---|---|
| PA 38:4 Me3 | 784,5 | 627,5 | 10 | 26 | 19 |
| PA 38:4 Me4 | 798,5 | 627,5 | 10 | 26 | 19 |
| PA 38:5 Me0 | 740,5 | 625,5 | 10 | 26 | 19 |
| PA 38:5 Me1 | 754,5 | 625,5 | 10 | 26 | 19 |
| PA 38:5 Me2 | 768,5 | 625,5 | 10 | 26 | 19 |
| PA 38:5 Me3 | 782,5 | 625,5 | 10 | 26 | 19 |
| PA 38:5 Me4 | 796,5 | 625,5 | 10 | 26 | 19 |
| PA 38:6 Me0 | 738,5 | 623,5 | 10 | 26 | 19 |
| PA 38:6 Me1 | 752,6 | 623,5 | 10 | 26 | 19 |
| PA 38:6 Me2 | 766,6 | 623,5 | 10 | 26 | 19 |
| PA 38:6 Me3 | 780,6 | 623,5 | 10 | 26 | 19 |
| PA 38:6 Me4 | 794,7 | 623,5 | 10 | 26 | 19 |
| PA 40:6 Me0 | 766,5 | 651,5 | 10 | 26 | 19 |
| PA 40:6 Me1 | 780,5 | 651,5 | 10 | 26 | 19 |
| PA 40:6 Me2 | 794,6 | 651,5 | 10 | 26 | 19 |
| PA 40:6 Me3 | 808,6 | 651,5 | 10 | 26 | 19 |
| PA 40:6 Me4 | 822,6 | 651,5 | 10 | 26 | 19 |
| PA 32:0 Me0 | 666,5 | 551,5 | 10 | 26 | 19 |
| PA 32:0 Me1 | 680,5 | 551,5 | 10 | 26 | 19 |
| PA 32:0 Me2 | 694,5 | 551,5 | 10 | 26 | 19 |
| PA 32:0 Me3 | 708,5 | 551,5 | 10 | 26 | 19 |
| PA 32:0 Me4 | 722,6 | 551,5 | 10 | 26 | 19 |
| PA 32:1 Me0 | 664,5 | 549,5 | 10 | 26 | 19 |
| PA 32:1 Me1 | 678,6 | 549,5 | 10 | 26 | 19 |
| PA 32:1 Me2 | 692,6 | 549,5 | 10 | 26 | 19 |
| PA 32:1 Me3 | 706,6 | 549,5 | 10 | 26 | 19 |
| PA 32:1 Me4 | 720,7 | 549,5 | 10 | 26 | 19 |
| PA 32:2 Me0 | 662,5 | 547,5 | 10 | 26 | 19 |
| PA 32:2 Me1 | 676,5 | 547,5 | 10 | 26 | 19 |
| PA 32:2 Me2 | 690,6 | 547,5 | 10 | 26 | 19 |
| PA 32:2 Me3 | 704,6 | 547,5 | 10 | 26 | 19 |
| PA 32:2 Me4 | 718,6 | 547,5 | 10 | 26 | 19 |
| PA 34:0 Me0 | 694,5 | 579,5 | 10 | 26 | 19 |
| PA 34:0 Me1 | 708,5 | 579,5 | 10 | 26 | 19 |
| PA 34:0 Me2 | 722,6 | 579,5 | 10 | 26 | 19 |
| PA 34:0 Me3 | 736,6 | 579,5 | 10 | 26 | 19 |
| PA 34:0 Me4 | 750,6 | 579,5 | 10 | 26 | 19 |
| PA 34:2 Me0 | 690,5 | 575,5 | 10 | 26 | 19 |
| PA 34:2 Me1 | 704,5 | 575,5 | 10 | 26 | 19 |
| PA 34:2 Me2 | 718,5 | 575,5 | 10 | 26 | 19 |
| PA 34:2 Me3 | 732,5 | 575,5 | 10 | 26 | 19 |
| PA 34:2 Me4 | 746,6 | 575,5 | 10 | 26 | 19 |
| PS 34:2 Me0 | 760,5 | 575,5 | 10 | 34 | 24 |
| PS 34:2 Me1 | 774,5 | 575,5 | 10 | 34 | 24 |
| PS 34:2 Me2 | 788,5 | 575,5 | 10 | 34 | 24 |
| PS 34:2 Me3 | 802,6 | 575,5 | 10 | 34 | 24 |
| PS 34:2 Me4 | 816,6 | 575,5 | 10 | 34 | 24 |
| PS 36:4 Me0 | 784,5 | 599,5 | 10 | 34 | 24 |
| PS 36:4 Me1 | 798,5 | 599,5 | 10 | 34 | 24 |
| PS 36:4 Me2 | 812,5 | 599,5 | 10 | 34 | 24 |
| PS 36:4 Me3 | 826,6 | 599,5 | 10 | 34 | 24 |
| PS 36:4 Me4 | 840,6 | 599,5 | 10 | 34 | 24 |
| PS 34:1 Me0 | 762,5 | 577,5 | 10 | 34 | 24 |
| PS 34:1 Me1 | 776,5 | 577,5 | 10 | 34 | 24 |
| PS 34:1 Me2 | 790,5 | 577,5 | 10 | 34 | 24 |
| PS 34:1 Me3 | 804,6 | 577,5 | 10 | 34 | 24 |
| PS 34:1 Me4 | 818,6 | 577,5 | 10 | 34 | 24 |
| PS 34:3 Me0 | 758,5 | 573,5 | 10 | 34 | 24 |
| PS 34:3 Me1 | 772,5 | 573,5 | 10 | 34 | 24 |
| PS 34:3 Me2 | 786,5 | 573,5 | 10 | 34 | 24 |
| PS 34:3 Me3 | 800,6 | 573,5 | 10 | 34 | 24 |
| PS 34:3 Me4 | 814,6 | 573,5 | 10 | 34 | 24 |
| PS 36:1 Me0 | 790,6 | 605,6 | 10 | 34 | 24 |

| | | | | | |
|---|---|---|---|---|---|
| PS 36:1 Me1 | 804,6 | 605,6 | 10 | 34 | 24 |
| PS 36:1 Me2 | 818,6 | 605,6 | 10 | 34 | 24 |
| PS 36:1 Me3 | 832,7 | 605,6 | 10 | 34 | 24 |
| PS 36:1 Me4 | 846,7 | 605,6 | 10 | 34 | 24 |
| PS 36:2 Me0 | 788,5 | 603,5 | 10 | 34 | 24 |
| PS 36:2 Me1 | 802,5 | 603,5 | 10 | 34 | 24 |
| PS 36:2 Me2 | 816,5 | 603,5 | 10 | 34 | 24 |
| PS 36:2 Me3 | 830,6 | 603,5 | 10 | 34 | 24 |
| PS 36:2 Me4 | 844,6 | 603,5 | 10 | 34 | 24 |
| PS 36:3 Me0 | 786,5 | 601,5 | 10 | 34 | 24 |
| PS 36:3 Me1 | 800,5 | 601,5 | 10 | 34 | 24 |
| PS 36:3 Me2 | 814,5 | 601,5 | 10 | 34 | 24 |
| PS 36:3 Me3 | 828,6 | 601,5 | 10 | 34 | 24 |
| PS 36:3 Me4 | 842,6 | 601,5 | 10 | 34 | 24 |
| PS 36:5 Me0 | 782,5 | 597,5 | 10 | 34 | 24 |
| PS 36:5 Me1 | 796,5 | 597,5 | 10 | 34 | 24 |
| PS 36:5 Me2 | 810,5 | 597,5 | 10 | 34 | 24 |
| PS 36:5 Me3 | 824,6 | 597,5 | 10 | 34 | 24 |
| PS 36:5 Me4 | 838,6 | 597,5 | 10 | 34 | 24 |
| PS 36:6 Me0 | 780,5 | 595,5 | 10 | 34 | 24 |
| PS 36:6 Me1 | 794,5 | 595,5 | 10 | 34 | 24 |
| PS 36:6 Me2 | 808,5 | 595,5 | 10 | 34 | 24 |
| PS 36:6 Me3 | 822,6 | 595,5 | 10 | 34 | 24 |
| PS 36:6 Me4 | 836,6 | 595,5 | 10 | 34 | 24 |
| PI4P 32:0 Me0 | 891,5 | 551,5 | 10 | 32 | 24 |
| PI4P 32:0 Me1 | 905,5 | 551,5 | 10 | 32 | 24 |
| PI4P 32:0 Me2 | 919,5 | 551,5 | 10 | 32 | 24 |
| PI4P 32:0 Me3 | 933,5 | 551,5 | 10 | 32 | 24 |
| PI4P 32:0 Me4 | 947,5 | 551,5 | 10 | 32 | 24 |
| PI4P 32:0 Me5 | 961,5 | 551,5 | 10 | 32 | 24 |
| PI4P 32:0 Me6 | 975,5 | 551,5 | 10 | 32 | 24 |
| PI4P 34:0 Me0 | 919,5 | 579,5 | 10 | 32 | 24 |
| PI4P 34:0 Me1 | 933,5 | 579,5 | 10 | 32 | 24 |
| PI4P 34:0 Me2 | 947,5 | 579,5 | 10 | 32 | 24 |
| PI4P 34:0 Me3 | 961,5 | 579,5 | 10 | 32 | 24 |
| PI4P 34:0 Me4 | 975,5 | 579,5 | 10 | 32 | 24 |
| PI4P 34:0 Me5 | 989,5 | 579,5 | 10 | 32 | 24 |
| PI4P 34:0 Me6 | 1003,5 | 579,5 | 10 | 32 | 24 |
| PI4P 34:1 Me0 | 917,5 | 577,5 | 10 | 32 | 24 |
| PI4P 34:1 Me1 | 931,5 | 577,5 | 10 | 32 | 24 |
| PI4P 34:1 Me2 | 945,5 | 577,5 | 10 | 32 | 24 |
| PI4P 34:1 Me3 | 959,5 | 577,5 | 10 | 32 | 24 |
| PI4P 34:1 Me4 | 973,5 | 577,5 | 10 | 32 | 24 |
| PI4P 34:1 Me5 | 987,5 | 577,5 | 10 | 32 | 24 |
| PI4P 34:1 Me6 | 1001,5 | 577,5 | 10 | 32 | 24 |
| PI4P 36:1 Me0 | 945,6 | 605,6 | 10 | 32 | 24 |
| PI4P 36:1 Me1 | 959,6 | 605,6 | 10 | 32 | 24 |
| PI4P 36:1 Me2 | 973,6 | 605,6 | 10 | 32 | 24 |
| PI4P 36:1 Me3 | 987,6 | 605,6 | 10 | 32 | 24 |
| PI4P 36:1 Me4 | 1001,6 | 605,6 | 10 | 32 | 24 |
| PI4P 36:1 Me5 | 1015,6 | 605,6 | 10 | 32 | 24 |
| PI4P 36:1 Me6 | 1029,6 | 605,6 | 10 | 32 | 24 |
| PI4P 36:2 Me0 | 943,5 | 603,5 | 10 | 32 | 24 |
| PI4P 36:2 Me1 | 957,5 | 603,5 | 10 | 32 | 24 |
| PI4P 36:2 Me2 | 971,5 | 603,5 | 10 | 32 | 24 |
| PI4P 36:2 Me3 | 985,5 | 603,5 | 10 | 32 | 24 |
| PI4P 36:2 Me4 | 999,5 | 603,5 | 10 | 32 | 24 |
| PI4P 36:2 Me5 | 1013,5 | 603,5 | 10 | 32 | 24 |
| PI4P 36:2 Me6 | 1027,5 | 603,5 | 10 | 32 | 24 |
| PI4P 36:3 Me0 | 941,5 | 601,5 | 10 | 32 | 24 |
| PI4P 36:3 Me1 | 955,5 | 601,5 | 10 | 32 | 24 |
| PI4P 36:3 Me2 | 969,5 | 601,5 | 10 | 32 | 24 |
| PI4P 36:3 Me3 | 983,5 | 601,5 | 10 | 32 | 24 |

| | | | | | |
|---|---|---|---|---|---|
| PI4P 36:3 Me4 | 997,5 | 601,5 | 10 | 32 | 24 |
| PI4P 36:3 Me5 | 1011,5 | 601,5 | 10 | 32 | 24 |
| PI4P 36:3 Me6 | 1025,5 | 601,5 | 10 | 32 | 24 |
| PI4P 36:4 Me0 | 939,5 | 599,5 | 10 | 32 | 24 |
| PI4P 36:4 Me1 | 953,5 | 599,5 | 10 | 32 | 24 |
| PI4P 36:4 Me2 | 967,5 | 599,5 | 10 | 32 | 24 |
| PI4P 36:4 Me3 | 981,5 | 599,5 | 10 | 32 | 24 |
| PI4P 36:4 Me4 | 995,5 | 599,5 | 10 | 32 | 24 |
| PI4P 36:4 Me5 | 1009,5 | 599,5 | 10 | 32 | 24 |
| PI4P 36:4 Me6 | 1023,5 | 599,5 | 10 | 32 | 24 |
| PI4P 38:2 Me0 | 971,6 | 631,6 | 10 | 32 | 24 |
| PI4P 38:2 Me1 | 985,6 | 631,6 | 10 | 32 | 24 |
| PI4P 38:2 Me2 | 999,6 | 631,6 | 10 | 32 | 24 |
| PI4P 38:2 Me3 | 1013,6 | 631,6 | 10 | 32 | 24 |
| PI4P 38:2 Me4 | 1027,6 | 631,6 | 10 | 32 | 24 |
| PI4P 38:2 Me5 | 1041,6 | 631,6 | 10 | 32 | 24 |
| PI4P 38:2 Me6 | 1055,6 | 631,6 | 10 | 32 | 24 |
| PI4P 38:3 Me0 | 969,6 | 629,6 | 10 | 32 | 24 |
| PI4P 38:3 Me1 | 983,6 | 629,6 | 10 | 32 | 24 |
| PI4P 38:3 Me2 | 997,6 | 629,6 | 10 | 32 | 24 |
| PI4P 38:3 Me3 | 1011,6 | 629,6 | 10 | 32 | 24 |
| PI4P 38:3 Me4 | 1025,6 | 629,6 | 10 | 32 | 24 |
| PI4P 38:3 Me5 | 1039,6 | 629,6 | 10 | 32 | 24 |
| PI4P 38:3 Me6 | 1053,6 | 629,6 | 10 | 32 | 24 |
| PI4P 38:4 Me0 | 967,5 | 627,5 | 10 | 32 | 24 |
| PI4P 38:4 Me1 | 981,5 | 627,5 | 10 | 32 | 24 |
| PI4P 38:4 Me2 | 995,5 | 627,5 | 10 | 32 | 24 |
| PI4P 38:4 Me3 | 1009,5 | 627,5 | 10 | 32 | 24 |
| PI4P 38:4 Me4 | 1023,5 | 627,5 | 10 | 32 | 24 |
| PI4P 38:4 Me5 | 1037,5 | 627,5 | 10 | 32 | 24 |
| PI4P 38:4 Me6 | 1051,5 | 627,5 | 10 | 32 | 24 |
| PI4P 38:5 Me0 | 965,5 | 625,5 | 10 | 32 | 24 |
| PI4P 38:5 Me1 | 979,5 | 625,5 | 10 | 32 | 24 |
| PI4P 38:5 Me2 | 993,5 | 625,5 | 10 | 32 | 24 |
| PI4P 38:5 Me3 | 1007,5 | 625,5 | 10 | 32 | 24 |
| PI4P 38:5 Me4 | 1021,5 | 625,5 | 10 | 32 | 24 |
| PI4P 38:5 Me5 | 1035,5 | 625,5 | 10 | 32 | 24 |
| PI4P 38:5 Me6 | 1049,5 | 625,5 | 10 | 32 | 24 |
| PI4P 40:4 Me0 | 995,6 | 655,6 | 10 | 32 | 24 |
| PI4P 40:4 Me1 | 1009,6 | 655,6 | 10 | 32 | 24 |
| PI4P 40:4 Me2 | 1023,6 | 655,6 | 10 | 32 | 24 |
| PI4P 40:4 Me3 | 1037,6 | 655,6 | 10 | 32 | 24 |
| PI4P 40:4 Me4 | 1051,6 | 655,6 | 10 | 32 | 24 |
| PI4P 40:4 Me5 | 1065,6 | 655,6 | 10 | 32 | 24 |
| PI4P 40:4 Me6 | 1079,6 | 655,6 | 10 | 32 | 24 |
| PI(4,5)$P_2$ 32:0 Me0 | 971,5 | 551,5 | 10 | 45 | 32 |
| PI(4,5)$P_2$ 32:0 Me1 | 985,5 | 551,5 | 10 | 45 | 32 |
| PI(4,5)$P_2$ 32:0 Me2 | 999,5 | 551,5 | 10 | 45 | 32 |
| PI(4,5)$P_2$ 32:0 Me3 | 1013,5 | 551,5 | 10 | 45 | 32 |
| PI(4,5)$P_2$ 32:0 Me4 | 1027,5 | 551,5 | 10 | 45 | 32 |
| PI(4,5)$P_2$ 32:0 Me5 | 1041,5 | 551,5 | 10 | 45 | 32 |
| PI(4,5)$P_2$ 32:0 Me6 | 1055,5 | 551,5 | 10 | 45 | 32 |
| PI(4,5)$P_2$ 32:0 Me7 | 1069,5 | 551,5 | 10 | 45 | 32 |
| PI(4,5)$P_2$ 34:0 Me0 | 999,5 | 579,5 | 10 | 45 | 32 |
| PI(4,5)$P_2$ 34:0 Me1 | 1013,5 | 579,5 | 10 | 45 | 32 |
| PI(4,5)$P_2$ 34:0 Me2 | 1027,5 | 579,5 | 10 | 45 | 32 |
| PI(4,5)$P_2$ 34:0 Me3 | 1041,5 | 579,5 | 10 | 45 | 32 |
| PI(4,5)$P_2$ 34:0 Me4 | 1055,5 | 579,5 | 10 | 45 | 32 |
| PI(4,5)$P_2$ 34:0 Me5 | 1069,5 | 579,5 | 10 | 45 | 32 |
| PI(4,5)$P_2$ 34:0 Me6 | 1083,5 | 579,5 | 10 | 45 | 32 |
| PI(4,5)$P_2$ 34:0 Me7 | 1097,5 | 579,5 | 10 | 45 | 32 |
| PI(4,5)$P_2$ 34:1 Me0 | 997,5 | 577,5 | 10 | 45 | 32 |
| PI(4,5)$P_2$ 34:1 Me1 | 1011,5 | 577,5 | 10 | 45 | 32 |

| | | | | | |
|---|---|---|---|---|---|
| PI(4,5)$P_2$ 34:1 Me2 | 1025,5 | 577,5 | 10 | 45 | 32 |
| PI(4,5)$P_2$ 34:1 Me3 | 1039,5 | 577,5 | 10 | 45 | 32 |
| PI(4,5)$P_2$ 34:1 Me4 | 1053,5 | 577,5 | 10 | 45 | 32 |
| PI(4,5)$P_2$ 34:1 Me5 | 1067,5 | 577,5 | 10 | 45 | 32 |
| PI(4,5)$P_2$ 34:1 Me6 | 1081,5 | 577,5 | 10 | 45 | 32 |
| PI(4,5)$P_2$ 34:1 Me7 | 1095,5 | 577,5 | 10 | 45 | 32 |
| PI(4,5)$P_2$ 36:1 Me0 | 1025,5 | 605,6 | 10 | 45 | 32 |
| PI(4,5)$P_2$ 36:1 Me1 | 1039,5 | 605,6 | 10 | 45 | 32 |
| PI(4,5)$P_2$ 36:1 Me2 | 1053,5 | 605,6 | 10 | 45 | 32 |
| PI(4,5)$P_2$ 36:1 Me3 | 1067,5 | 605,6 | 10 | 45 | 32 |
| PI(4,5)$P_2$ 36:1 Me4 | 1081,5 | 605,6 | 10 | 45 | 32 |
| PI(4,5)$P_2$ 36:1 Me5 | 1095,5 | 605,6 | 10 | 45 | 32 |
| PI(4,5)$P_2$ 36:1 Me6 | 1109,5 | 605,6 | 10 | 45 | 32 |
| PI(4,5)$P_2$ 36:1 Me7 | 1123,5 | 605,6 | 10 | 45 | 32 |
| PI(4,5)$P_2$ 36:2 Me0 | 1023,5 | 603,5 | 10 | 45 | 32 |
| PI(4,5)$P_2$ 36:2 Me1 | 1037,5 | 603,5 | 10 | 45 | 32 |
| PI(4,5)$P_2$ 36:2 Me2 | 1051,5 | 603,5 | 10 | 45 | 32 |
| PI(4,5)$P_2$ 36:2 Me3 | 1065,5 | 603,5 | 10 | 45 | 32 |
| PI(4,5)$P_2$ 36:2 Me4 | 1079,5 | 603,5 | 10 | 45 | 32 |
| PI(4,5)$P_2$ 36:2 Me5 | 1093,5 | 603,5 | 10 | 45 | 32 |
| PI(4,5)$P_2$ 36:2 Me6 | 1107,5 | 603,5 | 10 | 45 | 32 |
| PI(4,5)$P_2$ 36:2 Me7 | 1121,5 | 603,5 | 10 | 45 | 32 |
| PI(4,5)$P_2$ 36:3 Me0 | 1021,5 | 601,5 | 10 | 45 | 32 |
| PI(4,5)$P_2$ 36:3 Me1 | 1035,5 | 601,5 | 10 | 45 | 32 |
| PI(4,5)$P_2$ 36:3 Me2 | 1049,5 | 601,5 | 10 | 45 | 32 |
| PI(4,5)$P_2$ 36:3 Me3 | 1063,5 | 601,5 | 10 | 45 | 32 |
| PI(4,5)$P_2$ 36:3 Me4 | 1077,5 | 601,5 | 10 | 45 | 32 |
| PI(4,5)$P_2$ 36:3 Me5 | 1091,5 | 601,5 | 10 | 45 | 32 |
| PI(4,5)$P_2$ 36:3 Me6 | 1105,5 | 601,5 | 10 | 45 | 32 |
| PI(4,5)$P_2$ 36:3 Me7 | 1119,5 | 601,5 | 10 | 45 | 32 |
| PI(4,5)$P_2$ 36:4 Me0 | 1019,5 | 599,5 | 10 | 45 | 32 |
| PI(4,5)$P_2$ 36:4 Me1 | 1033,5 | 599,5 | 10 | 45 | 32 |
| PI(4,5)$P_2$ 36:4 Me2 | 1047,5 | 599,5 | 10 | 45 | 32 |
| PI(4,5)$P_2$ 36:4 Me3 | 1061,5 | 599,5 | 10 | 45 | 32 |
| PI(4,5)$P_2$ 36:4 Me4 | 1075,5 | 599,5 | 10 | 45 | 32 |
| PI(4,5)$P_2$ 36:4 Me5 | 1089,5 | 599,5 | 10 | 45 | 32 |
| PI(4,5)$P_2$ 36:4 Me6 | 1103,5 | 599,5 | 10 | 45 | 32 |
| PI(4,5)$P_2$ 36:4 Me7 | 1117,5 | 599,5 | 10 | 45 | 32 |
| PI(4,5)$P_2$ 38:3 Me0 | 1049,5 | 629,6 | 10 | 45 | 32 |
| PI(4,5)$P_2$ 38:3 Me1 | 1063,5 | 629,6 | 10 | 45 | 32 |
| PI(4,5)$P_2$ 38:3 Me2 | 1077,5 | 629,6 | 10 | 45 | 32 |
| PI(4,5)$P_2$ 38:3 Me3 | 1091,5 | 629,6 | 10 | 45 | 32 |
| PI(4,5)$P_2$ 38:3 Me4 | 1105,5 | 629,6 | 10 | 45 | 32 |
| PI(4,5)$P_2$ 38:3 Me5 | 1119,5 | 629,6 | 10 | 45 | 32 |
| PI(4,5)$P_2$ 38:3 Me6 | 1133,5 | 629,6 | 10 | 45 | 32 |
| PI(4,5)$P_2$ 38:3 Me7 | 1147,5 | 629,6 | 10 | 45 | 32 |
| PI(4,5)$P_2$ 38:4 Me0 | 1047,5 | 627,5 | 10 | 45 | 32 |
| PI(4,5)$P_2$ 38:4 Me1 | 1061,5 | 627,5 | 10 | 45 | 32 |
| PI(4,5)$P_2$ 38:4 Me2 | 1075,5 | 627,5 | 10 | 45 | 32 |
| PI(4,5)$P_2$ 38:4 Me3 | 1089,5 | 627,5 | 10 | 45 | 32 |
| PI(4,5)$P_2$ 38:4 Me4 | 1103,5 | 627,5 | 10 | 45 | 32 |
| PI(4,5)$P_2$ 38:4 Me5 | 1117,5 | 627,5 | 10 | 45 | 32 |
| PI(4,5)$P_2$ 38:4 Me6 | 1131,5 | 627,5 | 10 | 45 | 32 |
| PI(4,5)$P_2$ 38:4 Me7 | 1145,5 | 627,5 | 10 | 45 | 32 |
| PI(4,5)$P_2$ 38:5 Me0 | 1045,5 | 625,5 | 10 | 45 | 32 |
| PI(4,5)$P_2$ 38:5 Me1 | 1059,5 | 625,5 | 10 | 45 | 32 |
| PI(4,5)$P_2$ 38:5 Me2 | 1073,5 | 625,5 | 10 | 45 | 32 |
| PI(4,5)$P_2$ 38:5 Me3 | 1087,5 | 625,5 | 10 | 45 | 32 |
| PI(4,5)$P_2$ 38:5 Me4 | 1101,5 | 625,5 | 10 | 45 | 32 |
| PI(4,5)$P_2$ 38:5 Me5 | 1115,5 | 625,5 | 10 | 45 | 32 |
| PI(4,5)$P_2$ 38:5 Me6 | 1129,5 | 625,5 | 10 | 45 | 32 |
| PI(4,5)$P_2$ 38:5 Me7 | 1143,5 | 625,5 | 10 | 45 | 32 |
| PI(4,5)$P_2$ 40:4 Me0 | 1075,5 | 655,6 | 10 | 45 | 32 |

| | | | | | |
|---|---|---|---|---|---|
| PI(4,5)$P_2$ 40:4 Me1 | 1089,5 | 655,6 | 10 | 45 | 32 |
| PI(4,5)$P_2$ 40:4 Me2 | 1103,5 | 655,6 | 10 | 45 | 32 |
| PI(4,5)$P_2$ 40:4 Me3 | 1117,5 | 655,6 | 10 | 45 | 32 |
| PI(4,5)$P_2$ 40:4 Me4 | 1131,5 | 655,6 | 10 | 45 | 32 |
| PI(4,5)$P_2$ 40:4 Me5 | 1145,5 | 655,6 | 10 | 45 | 32 |
| PI(4,5)$P_2$ 40:4 Me6 | 1159,5 | 655,6 | 10 | 45 | 32 |
| PI(4,5)$P_2$ 40:4 Me7 | 1173,5 | 655,6 | 10 | 45 | 32 |
| PI(4,5)$P_2$ 40:5 Me0 | 1073,5 | 653,6 | 10 | 45 | 32 |
| PI(4,5)$P_2$ 40:5 Me1 | 1087,5 | 653,6 | 10 | 45 | 32 |
| PI(4,5)$P_2$ 40:5 Me2 | 1101,5 | 653,6 | 10 | 45 | 32 |
| PI(4,5)$P_2$ 40:5 Me3 | 1115,5 | 653,6 | 10 | 45 | 32 |
| PI(4,5)$P_2$ 40:5 Me4 | 1129,5 | 653,6 | 10 | 45 | 32 |
| PI(4,5)$P_2$ 40:5 Me5 | 1143,5 | 653,6 | 10 | 45 | 32 |
| PI(4,5)$P_2$ 40:5 Me6 | 1157,5 | 653,6 | 10 | 45 | 32 |
| PI(4,5)$P_2$ 40:5 Me7 | 1171,5 | 653,6 | 10 | 45 | 32 |

**Table S4.** List of MRM information (Q1 and Q3 mass, DP, CE and CXP) mass used for the confirmation of the analytical performance of the method

| Analyte | Q1 Mass (Da) | Q3 mass (Da) | DP (V) | CE (V) | CXP (V) |
|---|---|---|---|---|---|
| PA 18:0/18:1 Me2 | 748.6 | 605.6 | 10 | 26 | 19 |
| PA 18:0/18:1 Me0 | 720.6 | 605.6 | 10 | 26 | 19 |
| PS 16:0/18:1 Me2 | 790.6 | 577.5 | 10 | 34 | 24 |
| PS 16:0/18:1 Me0 | 762.5 | 577.5 | 10 | 34 | 24 |
| PS 18:0/18:2 Me2 | 816.6 | 603.5 | 10 | 34 | 24 |
| PS 18:0/18:2 Me0 | 788.5 | 603.5 | 10 | 34 | 24 |
| PI 16:0/18:1 Me1 | 868.6 | 577.5 | 10 | 31 | 25 |
| PI 16:0/18:1 Me0 | 854.6 | 577.5 | 10 | 31 | 25 |
| PI 17:0/14:1 Me1 | 826.6 | 535.5 | 10 | 31 | 25 |
| PI 17:0/14:1 Me0 | 812.5 | 535.5 | 10 | 31 | 25 |
| PI4P 17:0/20:4 Me3 | 995.6 | 613.5 | 10 | 32 | 24 |
| PI4P 17:0/20:4 Me0 | 953.5 | 613.5 | 10 | 32 | 24 |
| PI(4,5)$P_2$ 17:0/20:4 Me5 | 1103.6 | 613.5 | 10 | 45 | 32 |
| PI(4,5)$P_2$ 17:0/20:4 Me0 | 1033.5 | 613.5 | 10 | 45 | 32 |
| PS 17:0/17:0 Me2 (IS) | 792.6 | 579.5 | 10 | 34 | 24 |

**Table S5.** List of MRM information (Q1 and Q3 mass, DP, CE and CXP) used for the analysis of plant samples

| Analyte | Q1 Mass (Da) | Q3 mass (Da) | DP (V) | CE (V) | CXP (V) |
|---|---|---|---|---|---|
| PA 32:0 | 694.5 | 551.5 | 10 | 26 | 19 |
| PA 32:1 | 692.5 | 549.5 | 10 | 26 | 19 |
| PA 32:2 | 690.5 | 547.5 | 10 | 26 | 19 |
| PA 34:0 | 722.6 | 579.5 | 10 | 26 | 19 |
| PA 34:1 | 720.6 | 577.5 | 10 | 26 | 19 |
| PA 34:2 | 718.5 | 575.5 | 10 | 26 | 19 |
| PA 34:3 | 716.5 | 573.5 | 10 | 26 | 19 |
| PA 34:4 | 714.5 | 571.5 | 10 | 26 | 19 |
| PA 36:0 | 750.6 | 607.6 | 10 | 26 | 19 |
| PA 36:1 | 748.6 | 605.6 | 10 | 26 | 19 |
| PA 36:2 | 746.6 | 603.5 | 10 | 26 | 19 |
| PA 36:3 | 744.6 | 601.5 | 10 | 26 | 19 |
| PA 36:4 | 742.5 | 599.5 | 10 | 26 | 19 |
| PA 36:5 | 740.5 | 597.5 | 10 | 26 | 19 |
| PA 36:6 | 738.5 | 595.5 | 10 | 26 | 19 |
| PA 38:0 | 778.6 | 635.6 | 10 | 26 | 19 |
| PA 38:1 | 776.6 | 633.6 | 10 | 26 | 19 |
| PA 38:2 | 774.6 | 631.6 | 10 | 26 | 19 |
| PA 38:3 | 772.6 | 629.6 | 10 | 26 | 19 |
| PA 38:4 | 770.6 | 627.5 | 10 | 26 | 19 |
| PA 40:0 | 806.7 | 663.6 | 10 | 26 | 19 |
| PA 40:1 | 804.7 | 661.6 | 10 | 26 | 19 |
| PA 40:2 | 802.6 | 659.6 | 10 | 26 | 19 |
| PA 40:3 | 800.6 | 657.6 | 10 | 26 | 19 |
| PA 40:4 | 798.6 | 655.6 | 10 | 26 | 19 |
| PA 42:0 | 834.7 | 691.7 | 10 | 26 | 19 |
| PA 42:1 | 832.7 | 689.6 | 10 | 26 | 19 |
| PA 42:2 | 830.7 | 687.6 | 10 | 26 | 19 |
| PA 42:3 | 828.7 | 685.6 | 10 | 26 | 19 |
| PA 42:4 | 826.6 | 683.6 | 10 | 26 | 19 |
| PA 44:0 | 862.7 | 719.7 | 10 | 26 | 19 |
| PA 44:1 | 860.7 | 717.7 | 10 | 26 | 19 |
| PA 44:2 | 858.7 | 715.7 | 10 | 26 | 19 |
| PA 44:3 | 856.7 | 713.6 | 10 | 26 | 19 |
| PA 44:4 | 854.7 | 711.6 | 10 | 26 | 19 |
| PA 46:0 | 890.8 | 747.7 | 10 | 26 | 19 |
| PA 46:1 | 888.8 | 745.7 | 10 | 26 | 19 |
| PA 46:2 | 886.7 | 743.7 | 10 | 26 | 19 |
| PA 48:0 | 918.8 | 775.8 | 10 | 26 | 19 |
| PA 48:1 | 916.8 | 773.7 | 10 | 26 | 19 |
| PA 48:2 | 914.8 | 771.7 | 10 | 26 | 19 |
| PA 50:0 | 946.8 | 803.8 | 10 | 26 | 19 |
| PA 50:1 | 944.8 | 801.8 | 10 | 26 | 19 |
| PA 50:2 | 942.8 | 799.8 | 10 | 26 | 19 |
| PA 52:0 | 974.9 | 831.8 | 10 | 26 | 19 |
| PA 52:1 | 972.8 | 829.8 | 10 | 26 | 19 |
| PA 52:2 | 970.8 | 827.8 | 10 | 26 | 19 |
| PA 33:1 (IS) | 706.5 | 563.5 | 10 | 26 | 19 |
| PI 32:0 | 842.6 | 551.5 | 10 | 31 | 25 |
| PI 32:1 | 840.6 | 549.5 | 10 | 31 | 25 |
| PI 32:2 | 838.5 | 547.5 | 10 | 31 | 25 |
| PI 34:0 | 870.6 | 579.5 | 10 | 31 | 25 |
| PI 34:1 | 868.6 | 577.5 | 10 | 31 | 25 |
| PI 34:2 | 866.6 | 575.5 | 10 | 31 | 25 |
| PI 34:3 | 864.6 | 573.5 | 10 | 31 | 25 |
| PI 34:4 | 862.5 | 571.5 | 10 | 31 | 25 |
| PI 36:0 | 898.6 | 607.6 | 10 | 31 | 25 |

| | | | | | |
|---|---|---|---|---|---|
| PI 36:1 | 896.6 | 605.6 | 10 | 31 | 25 |
| PI 36:2 | 894.6 | 603.5 | 10 | 31 | 25 |
| PI 36:3 | 892.6 | 601.5 | 10 | 31 | 25 |
| PI 36:4 | 890.6 | 599.5 | 10 | 31 | 25 |
| PI 36:5 | 888.6 | 597.5 | 10 | 31 | 25 |
| PI 36:6 | 886.5 | 595.5 | 10 | 31 | 25 |
| PI 38:0 | 926.7 | 635.6 | 10 | 31 | 25 |
| PI 38:1 | 924.7 | 633.6 | 10 | 31 | 25 |
| PI 38:2 | 922.6 | 631.6 | 10 | 31 | 25 |
| PI 38:3 | 920.6 | 629.6 | 10 | 31 | 25 |
| PI 38:4 | 918.6 | 627.5 | 10 | 31 | 25 |
| PI 40:0 | 954.7 | 663.6 | 10 | 31 | 25 |
| PI 40:1 | 952.7 | 661.6 | 10 | 31 | 25 |
| PI 40:2 | 950.7 | 659.6 | 10 | 31 | 25 |
| PI 42:0 | 982.7 | 691.7 | 10 | 31 | 25 |
| PI 42:1 | 980.7 | 689.6 | 10 | 31 | 25 |
| PI 42:2 | 978.7 | 687.6 | 10 | 31 | 25 |
| PI 42:3 | 976.7 | 685.6 | 10 | 31 | 25 |
| PI 42:4 | 974.7 | 683.6 | 10 | 31 | 25 |
| PI 44:0 | 1010.8 | 719.7 | 10 | 31 | 25 |
| PI 44:1 | 1008.8 | 717.7 | 10 | 31 | 25 |
| PI 44:2 | 1006.7 | 715.7 | 10 | 31 | 25 |
| PI 44:3 | 1004.7 | 713.6 | 10 | 31 | 25 |
| PI 44:4 | 1002.7 | 711.6 | 10 | 31 | 25 |
| PI 46:0 | 1038.8 | 747.7 | 10 | 31 | 25 |
| PI 46:1 | 1036.8 | 745.7 | 10 | 31 | 25 |
| PI 46:2 | 1034.8 | 743.7 | 10 | 31 | 25 |
| PI 48:0 | 1066.8 | 775.8 | 10 | 31 | 25 |
| PI 48:1 | 1064.8 | 773.7 | 10 | 31 | 25 |
| PI 48:2 | 1062.8 | 771.7 | 10 | 31 | 25 |
| PI 50:0 | 1094.9 | 803.8 | 10 | 31 | 25 |
| PI 50:1 | 1092.8 | 801.8 | 10 | 31 | 25 |
| PI 50:2 | 1090.8 | 799.8 | 10 | 31 | 25 |
| PI 52:0 | 1122.9 | 831.8 | 10 | 31 | 25 |
| PI 52:1 | 1120.9 | 829.8 | 10 | 31 | 25 |
| PI 52:2 | 1118.9 | 827.8 | 10 | 31 | 25 |
| PS 32:0 | 764.6 | 551.5 | 10 | 34 | 24 |
| PS 32:1 | 762.5 | 549.5 | 10 | 34 | 24 |
| PS 32:2 | 760.5 | 547.5 | 10 | 34 | 24 |
| PS 34:0 | 792.6 | 579.5 | 10 | 34 | 24 |
| PS 34:1 | 790.6 | 577.5 | 10 | 34 | 24 |
| PS 34:2 | 788.6 | 575.5 | 10 | 34 | 24 |
| PS 34:3 | 786.5 | 573.5 | 10 | 34 | 24 |
| PS 34:4 | 784.5 | 571.5 | 10 | 34 | 24 |
| PS 36:0 | 820.6 | 607.6 | 10 | 34 | 24 |
| PS 36:1 | 818.6 | 605.6 | 10 | 34 | 24 |
| PS 36:2 | 816.6 | 603.5 | 10 | 34 | 24 |
| PS 36:3 | 814.6 | 601.5 | 10 | 34 | 24 |
| PS 36:4 | 812.6 | 599.5 | 10 | 34 | 24 |
| PS 36:5 | 810.5 | 597.5 | 10 | 34 | 24 |
| PS 36:6 | 808.5 | 595.5 | 10 | 34 | 24 |
| PS 38:0 | 848.6 | 635.6 | 10 | 34 | 24 |
| PS 38:1 | 846.6 | 633.6 | 10 | 34 | 24 |
| PS 38:2 | 844.6 | 631.6 | 10 | 34 | 24 |
| PS 38:3 | 842.6 | 629.6 | 10 | 34 | 24 |
| PS 38:4 | 840.6 | 627.5 | 10 | 34 | 24 |
| PS 40:0 | 876.7 | 663.6 | 10 | 34 | 24 |
| PS 40:1 | 874.7 | 661.6 | 10 | 34 | 24 |
| PS 40:2 | 872.6 | 659.6 | 10 | 34 | 24 |
| PS 40:3 | 870.6 | 657.6 | 10 | 34 | 24 |
| PS 40:4 | 868.6 | 655.6 | 10 | 34 | 24 |
| PS 42:0 | 904.7 | 691.7 | 10 | 34 | 24 |
| PS 42:1 | 902.7 | 689.6 | 10 | 34 | 24 |

| | | | | | |
|---|---|---|---|---|---|
| PS 42:2 | 900.7 | 687.6 | 10 | 34 | 24 |
| PS 42:3 | 898.7 | 685.6 | 10 | 34 | 24 |
| PS 42:4 | 896.6 | 683.6 | 10 | 34 | 24 |
| PS 44:0 | 932.7 | 719.7 | 10 | 34 | 24 |
| PS 44:1 | 930.7 | 717.7 | 10 | 34 | 24 |
| PS 44:2 | 928.7 | 715.7 | 10 | 34 | 24 |
| PS 44:3 | 926.7 | 713.6 | 10 | 34 | 24 |
| PS 44:4 | 924.7 | 711.6 | 10 | 34 | 24 |
| PS 46:0 | 960.8 | 747.7 | 10 | 34 | 24 |
| PS 46:1 | 958.8 | 745.7 | 10 | 34 | 24 |
| PS 46:2 | 956.7 | 743.7 | 10 | 34 | 24 |
| PS 48:0 | 988.8 | 775.8 | 10 | 34 | 24 |
| PS 48:1 | 986.8 | 773.7 | 10 | 34 | 24 |
| PS 48:2 | 984.8 | 771.7 | 10 | 34 | 24 |
| PS 50:0 | 1016.8 | 803.8 | 10 | 34 | 24 |
| PS 50:1 | 1014.8 | 801.8 | 10 | 34 | 24 |
| PS 50:2 | 1012.8 | 799.8 | 10 | 34 | 24 |
| PS 52:0 | 1044.9 | 831.8 | 10 | 34 | 24 |
| PS 52:1 | 1042.8 | 829.8 | 10 | 34 | 24 |
| PS 52:2 | 1040.8 | 827.8 | 10 | 34 | 24 |
| PIP 32:0 | 933.6 | 551.5 | 10 | 32 | 24 |
| PIP 32:1 | 931.5 | 549.5 | 10 | 32 | 24 |
| PIP 32:2 | 929.5 | 547.5 | 10 | 32 | 24 |
| PIP 34:0 | 961.6 | 579.5 | 10 | 32 | 24 |
| PIP 34:1 | 959.6 | 577.5 | 10 | 32 | 24 |
| PIP 34:2 | 957.6 | 575.5 | 10 | 32 | 24 |
| PIP 34:3 | 955.5 | 573.5 | 10 | 32 | 24 |
| PIP 34:4 | 953.5 | 571.5 | 10 | 32 | 24 |
| PIP 36:0 | 989.6 | 607.6 | 10 | 32 | 24 |
| PIP 36:1 | 987.6 | 605.6 | 10 | 32 | 24 |
| PIP 36:2 | 985.6 | 603.5 | 10 | 32 | 24 |
| PIP 36:3 | 983.6 | 601.5 | 10 | 32 | 24 |
| PIP 36:4 | 981.6 | 599.5 | 10 | 32 | 24 |
| PIP 36:5 | 979.5 | 597.5 | 10 | 32 | 24 |
| PIP 36:6 | 977.5 | 595.5 | 10 | 32 | 24 |
| PIP 38:0 | 1017.7 | 635.6 | 10 | 32 | 24 |
| PIP 38:1 | 1015.6 | 633.6 | 10 | 32 | 24 |
| PIP 38:2 | 1013.6 | 631.6 | 10 | 32 | 24 |
| PIP 38:3 | 1011.6 | 629.6 | 10 | 32 | 24 |
| PIP 38:4 | 1009.6 | 627.5 | 10 | 32 | 24 |
| PIP 40:0 | 1045.7 | 663.6 | 10 | 32 | 24 |
| PIP 40:1 | 1043.7 | 661.6 | 10 | 32 | 24 |
| PIP 40:2 | 1041.7 | 659.6 | 10 | 32 | 24 |
| PIP 40:3 | 1039.6 | 657.6 | 10 | 32 | 24 |
| PIP 40:4 | 1037.6 | 655.6 | 10 | 32 | 24 |
| PIP 42:0 | 1073.7 | 691.7 | 10 | 32 | 24 |
| PIP 42:1 | 1071.7 | 689.6 | 10 | 32 | 24 |
| PIP 42:2 | 1069.7 | 687.6 | 10 | 32 | 24 |
| PIP 42:3 | 1067.7 | 685.6 | 10 | 32 | 24 |
| PIP 42:4 | 1065.7 | 683.6 | 10 | 32 | 24 |
| PIP 44:0 | 1101.7 | 719.7 | 10 | 32 | 24 |
| PIP 44:1 | 1099.7 | 717.7 | 10 | 32 | 24 |
| PIP 44:2 | 1097.7 | 715.7 | 10 | 32 | 24 |
| PIP 44:3 | 1095.7 | 713.6 | 10 | 32 | 24 |
| PIP 44:4 | 1093.7 | 711.6 | 10 | 32 | 24 |
| PIP 46:0 | 1129.8 | 747.7 | 10 | 32 | 24 |
| PIP 46:1 | 1127.8 | 745.7 | 10 | 32 | 24 |
| PIP 46:2 | 1125.7 | 743.7 | 10 | 32 | 24 |
| PIP 48:0 | 1157.8 | 775.8 | 10 | 32 | 24 |
| PIP 48:1 | 1155.8 | 773.7 | 10 | 32 | 24 |
| PIP 48:2 | 1153.8 | 771.7 | 10 | 32 | 24 |
| PIP 50:0 | 1185.8 | 803.8 | 10 | 32 | 24 |
| PIP 50:1 | 1183.8 | 801.8 | 10 | 32 | 24 |

| | | | | | |
|---|---|---|---|---|---|
| PIP 50:2 | 1181.8 | 799.8 | 10 | 32 | 24 |
| PIP 52:0 | 1213.9 | 831.8 | 10 | 32 | 24 |
| PIP 52:1 | 1211.9 | 829.8 | 10 | 32 | 24 |
| PIP 52:2 | 1209.8 | 827.8 | 10 | 32 | 24 |
| $PIP_2$ 32:0 | 1041.6 | 551.5 | 10 | 45 | 32 |
| $PIP_2$ 32:1 | 1039.6 | 549.5 | 10 | 45 | 32 |
| $PIP_2$ 32:2 | 1037.5 | 547.5 | 10 | 45 | 32 |
| $PIP_2$ 34:0 | 1069.6 | 579.5 | 10 | 45 | 32 |
| $PIP_2$ 34:1 | 1067.6 | 577.5 | 10 | 45 | 32 |
| $PIP_2$ 34:2 | 1065.6 | 575.5 | 10 | 45 | 32 |
| $PIP_2$ 34:3 | 1063.6 | 573.5 | 10 | 45 | 32 |
| $PIP_2$ 34:4 | 1061.5 | 571.5 | 10 | 45 | 32 |
| $PIP_2$ 36:0 | 1097.6 | 607.6 | 10 | 45 | 32 |
| $PIP_2$ 36:1 | 1095.6 | 605.6 | 10 | 45 | 32 |
| $PIP_2$ 36:2 | 1093.6 | 603.5 | 10 | 45 | 32 |
| $PIP_2$ 36:3 | 1091.6 | 601.5 | 10 | 45 | 32 |
| $PIP_2$ 36:4 | 1089.6 | 599.5 | 10 | 45 | 32 |
| $PIP_2$ 36:5 | 1087.6 | 597.5 | 10 | 45 | 32 |
| $PIP_2$ 36:6 | 1085.5 | 595.5 | 10 | 45 | 32 |
| $PIP_2$ 38:0 | 1125.7 | 635.6 | 10 | 45 | 32 |
| $PIP_2$ 38:1 | 1123.6 | 633.6 | 10 | 45 | 32 |
| $PIP_2$ 38:2 | 1121.6 | 631.6 | 10 | 45 | 32 |
| $PIP_2$ 38:3 | 1119.6 | 629.6 | 10 | 45 | 32 |
| $PIP_2$ 38:4 | 1117.6 | 627.5 | 10 | 45 | 32 |
| $PIP_2$ 40:0 | 1153.7 | 663.6 | 10 | 45 | 32 |
| $PIP_2$ 40:1 | 1151.7 | 661.6 | 10 | 45 | 32 |
| $PIP_2$ 40:2 | 1149.7 | 659.6 | 10 | 45 | 32 |
| $PIP_2$ 40:3 | 1147.6 | 657.6 | 10 | 45 | 32 |
| $PIP_2$ 40:4 | 1145.6 | 655.6 | 10 | 45 | 32 |
| $PIP_2$ 42:0 | 1181.7 | 691.7 | 10 | 45 | 32 |
| $PIP_2$ 42:1 | 1179.7 | 689.6 | 10 | 45 | 32 |
| $PIP_2$ 42:2 | 1177.7 | 687.6 | 10 | 45 | 32 |
| $PIP_2$ 42:3 | 1175.7 | 685.6 | 10 | 45 | 32 |
| $PIP_2$ 42:4 | 1173.7 | 683.6 | 10 | 45 | 32 |
| $PIP_2$ 44:0 | 1209.8 | 719.7 | 10 | 45 | 32 |
| $PIP_2$ 44:1 | 1207.7 | 717.7 | 10 | 45 | 32 |
| $PIP_2$ 44:2 | 1205.7 | 715.7 | 10 | 45 | 32 |
| $PIP_2$ 44:3 | 1203.7 | 713.6 | 10 | 45 | 32 |
| $PIP_2$ 44:4 | 1201.7 | 711.6 | 10 | 45 | 32 |
| $PIP_2$ 46:0 | 1237.8 | 747.7 | 10 | 45 | 32 |
| $PIP_2$ 46:1 | 1235.8 | 745.7 | 10 | 45 | 32 |
| $PIP_2$ 46:2 | 1233.8 | 743.7 | 10 | 45 | 32 |
